# Leaf and shoot apical meristem transcriptomes of quinoa (*Chenopodium quinoa* Willd.) in response to photoperiod and plant development

**DOI:** 10.1101/2023.08.31.555728

**Authors:** Nathaly Maldonado-Taipe, Elodie Rey, Mark Tester, Christian Jung, Nazgol Emrani

## Abstract

Our study aimed to identify candidate genes for flowering time regulation and photoperiod response in quinoa. We investigated the timing of photoperiod-driven floral transition and analyzed the transcriptomes of leaf and shoot apical meristems in photoperiod-sensitive and -insensitive quinoa accessions. Histology analysis of the apical meristem showed that floral transition in quinoa initiates two to three weeks after sowing. We found four groups of differentially expressed genes responding to plant development and floral transition, which were annotated in the QQ74-V2 reference genome, including (i) 222 genes differentially responding to photoperiod in leaves, (ii) 1,812 genes differentially expressed between accessions under long-day conditions in leaves, (iii) 57 genes responding to developmental changes between weeks under short-day conditions in leaves, and (iv) 911 genes responding to floral transition within the shoot apical meristem. Interestingly, out of the thousands of candidates, two putative *FT* orthologues and several others have been reported as key regulators of flowering time in other species (e.g., *SOC1*, *COL*, *AP1*). Additionally, we used co-expression networks to associate novel transcripts to a putative biological process based on the annotated genes within the same co-expression cluster. The candidate genes in this study would benefit quinoa breeding by identifying and integrating their beneficial haplotypes in crossing programs to develop adapted cultivars to diverse environmental conditions.

## Introduction

The pseudocereal quinoa (*Chenopodium quinoa* Willd.) is an allotetraploid species (2*n* = 4*x* = 36) with a genome size of 1.45-1.50 Gb that encompasses 44,776 genes (Jarvis *et al*., 2017). It resulted from a hybridization event between an A-genome diploid species that most likely originated from a relative of *C. pallidicaule,* and a B-genome diploid species (*C. suecicum, C. ficifolium* or another related diploid species) (Štorchová *et al*., 2015; Jarvis *et al*., 2017). Quinoa originated from the Andean region of South America, where it has grown for over 6,000 years as a subsistence crop. Quinoa seeds have a high protein content and an exceptional amino acid composition (Iqbal *et al*., 2020; Granado-Rodríguez *et al*., 2021). Moreover, quinoa is tolerant against frost, drought, and salinity (Kiani-Pouya *et al*., 2022). Because of these advantages, it has gained substantial attention as a human diet, and breeding and cultivation has been initiated in over 120 countries (Alandia *et al*., 2020).

The short-day nature of this crop remains an obstacle to its cultivation in temperate regions and high latitudes in Europe, North America, and China (Murphy *et al*., 2018; Patiranage *et al*., 2021). A better understanding of the genetic mechanisms underlying floral transition, including photoperiodic regulation of flowering time, will help address this problem. Furthermore, breeding programs aiming to adapt quinoa to new environments can benefit from the profound effect of flowering time control on crop adaptation and yield potential (Gaudinier & Blackman, 2020; Patiranage *et al*., 2022).

Fuller (1949) published the first report on quinoa’s response to photoperiod, where short photoperiods induced earlier inflorescence appearance and flowering than long photoperiods. Since then, only a few studies have elaborated on quinoa’s genetic mechanisms of flowering time regulation. Golicz *et al*. (2020) used a computational approach to identify hundreds of putative orthologues of *Arabidopsis thaliana* flowering time genes in quinoa. They found 611 genes with high sequence homology to *A. thaliana* flowering genes, which could be considered putative flowering time regulators in quinoa. In another study, long non-coding RNA (lncRNA) was analyzed under short-day (SD) conditions with a 60 min night-break (NB) (Wu *et al*., 2021). The authors identified 24 lncRNA involved in flowering time regulation, some of which targeted *FLOWERING LOCUS T* (*FT*) and *TWIN SISTER of FT* (*TSF*) homologs (based on *in silico* analysis). Moreover, they found that quinoa homologs of *CONSTANS-like* (*COL*), *LATE ELONGATED HYPOCOTYL* (*LHY*)*, EARLY FLOWERING 3* (*ELF3*), and *ELONGATED HYPOCOTYL 5* (*HY5*) were down-regulated after NB. In contrast, *PHYTOCROME A (PHYA)* and *CRYPTOCHROME1 (CRY1)* homologs were up-regulated after NB. Furthermore, the role of *FT* and *COL* homologs in adapting quinoa to different day-length conditions was highlighted by Patiranage *et al*. (2021). In their study, the haplotypes of 12 putative flowering time genes were analyzed using a set of 276 accessions grown under long-day (LD) and SD conditions. As a result, *CqFT1A*, *CqCOL2B*, *CqCOL4A-1*, and *CqCOL5B* were associated with flowering time variation under LD but not under SD conditions. Recently, a transcriptome study of diurnally collected quinoa samples unveiled *CO-like* transcription factors among the diurnally regulated genes sensitive to the switch from LD to SD (Wu *et al*., 2023).

The regulation of flowering time has also been studied in quinoa-closely-related species. In *Chenopodium rubrum*, *FLOWERING LOCUS T-LIKE 1* (*CrFTL1*) change of expression was associated with experimental conditions which led to flowering, whereas *CrFTL2* was constitutively expressed (Cháb *et al*., 2008). Moreover, two *CrCOL* genes were down-regulated during the light period regardless of the length of the preceding dark period. Likewise, the floral promoter *CrFTL1* was down-regulated during the light period (Drabešová *et al*., 2014). In another study, using a segregating *Chenopodium ficifolium* F_2_ population, sequence variations at an *FTL1* ortholog explained flowering time, plant height, and branching. In a recent study, the transcriptome of *C. ficifolium* was studied at four different developmental stages under LD and SD conditions. The authors identified 6,096 differentially expressed genes (DEGs) possibly associated with floral induction. The most relevant candidate genes were those responsible for phytohormone metabolism and signaling since enhanced cytokinin content and the stimulation of cytokinin and gibberellic acid signaling pathways correlated with floral induction under short days (Gutierrez-Larruscain *et al*., 2022).

This study aimed to determine when the shoot apical meristem (SAM) is turned into a floral meristem. Furthermore, we targeted genes controlling flowering time and photoperiod response in quinoa. We expected those genes to display differential expression profiles between photoperiods and accessions with contrasting life cycle regimes. Accordingly, we analyzed the development of the SAM in two accessions differentially responding to day length. Studying leaf and SAM transcriptomes from plants grown under SD and LD conditions resulted in thousands of differentially expressed genes. Our study provides new insight into quinoa’s flowering time and photoperiod regulation.

## Materials and Methods

### Plant material and growth conditions

Two quinoa accessions were investigated in this study, D-12082 from Peru (seed code: 182301) and PI-614886 from Chile (the sequenced quinoa reference genome QQ74 (Jarvis *et al*., 2017), seed code: 182283), which differentially flowered under SD and LD conditions as reported by Patiranage *et al*. (2021). Plants were grown under two day-length regimes (LD and SD). One hundred five plants of each accession were grown in 3 × 3 cm 35-multiwell palettes (Hermann Meyer KG, Germany) in a growth chamber under long-day conditions (LD) (22 °C and 16 h light; 900 μmol·m^−2^·s^−1^, Son-T Agro 400 W, Koninklijke Philips Electronics N.V., Eindhoven, The Netherlands). Likewise, 105 plants of each accession were grown under short-day conditions (SD) (22 °C and 8 h light; 900 μmol·m^−2^·s^−1^).

### Phenotyping and histological analysis

In each experiment (LD and SD), we phenotyped ten plants per accession for days to bolting (days until the floral bud is visible) and days to flowering (days until the first flower opens) (Stanschewski *et al*., 2021). We sampled seven apices per accession and growth conditions weekly until plants reached the bolting stage (Supplementary Table 1). We used these apices for histological analysis. Apices were fixed in 4.0% FAA (4.0% formaldehyde, 50.0% ethanol, and 5.0% acetic acid) overnight. Then, the samples were dehydrated by an ethanol series and embedded in Paraplast (Sigma, P3683) by a standard protocol (Wu & Wagner, 2012). Apices were sectioned at 8 μm, using a rotatory microtome (Leica RM 2255, Wetzlar, Germany) and stained with 0.05% toluidine blue.

### RNA isolation and DNAse treatment

Leaves were harvested at ZT-9 (Zeitgeber-time) and SAMs at ZT-9 to ZT-12 from two accessions at different developmental stages under LD and SD conditions (Supplementary Table 1). Samples were immediately placed in liquid nitrogen. RNA was isolated from 48 leaf and 24 SAM samples using the peqGold Total RNA Kit (PeqLab, Erlangen, Germany) protocol. Ten to 14 SAMs were pooled to make one biological replicate. After RNA isolation, we performed a DNase I treatment (Thermo Fisher Scientific Inc., Waltham, United States) for 30 min at 37°C on the isolated RNA to eliminate DNA contamination.

### Sequencing, reads alignment, and transcript assembly

Directional mRNA library preparation was carried out by the Poly(A) enrichment method at Novogene Company Limited (Cambridge, United Kingdom). Seventy-two cDNA libraries, prepared from the DNase-treated RNA samples (RIN number >6.9), were sequenced in 2×150 bp paired-rend (PE) using NovaSeq 6000 PE150.

We obtained read-count data from the raw reads following the protocol described by Pertea *et al*. (2016). Briefly, reads from each sample were first mapped to the QQ74-V2 reference genome of *C. quinoa* cv. QQ74 (CoGe Genome ID: id60716) with HISAT2 (v.2.1.9) (Kim *et al*., 2015). Then, novel and known transcripts were assembled for each sample and merged using StringTie (v.2.2.0) (Pertea *et al*., 2015). The transcriptome was then retrieved from the merged annotation using the gffread function of GFF utilities (v.0.11.1) (http://github.com/gpertea/gffcompare) (Pertea & Pertea, 2020). Next, we obtained the read counts by mapping reads back to the transcripts using bowtie2 (v.2.3.5) (Langmead & Salzberg, 2012) leveraged by rsem (v.1.3.1) (Li & Dewey, 2011) with ‘very_sensitive’ stringency level. Finally, we extracted the ‘expected counts’ from each sample to build the gene matrix containing known and novel transcripts. We termed sequences as “novel transcripts” when they were assembled as transcripts by StringTie (v.2.2.0) but were not annotated in the reference genome.

### Differential expression analysis

The differential expression analysis was carried out at the gene level using R (version 4.1.1) package edgeR (version 3.36.0) (Chen, Y *et al*., 2020). First, we excluded lowly expressed genes by keeping genes with about five read counts or more in a minimum number of samples, where the number of samples is chosen according to the minimum group sample size. This filtering uses CPM values rather than counts to avoid giving preference to samples with large library sizes. Later, we normalized the gene expression by the ‘Trimmed mean of M-values normalization method’ (TMM) (Robinson & Oshlack, 2010), which estimates scale factors between samples, to take library composition into account; thus, obtaining TMM normalized counts per million (CPM). To view detailed gene expression trends among samples, the Trimmed Mean log ratios (log_2_ fold change of M values) (Robinson & Oshlack, 2010) were used for multidimensional scaling (MDS) plots. Next, we used the dgeFitGLM function to fit a generalized linear model to our count data and conducted gene-wise statistical tests for our contrasts. We defined a differentially expressed gene (DEG) as the one with a false discovery rate < 0.05 and a log_2_ fold change (FC) ≥ 0.25 or ≤ −0.25.

We investigated the DEGs in two main analyses. First, we used the DEGs of leaf and SAM transcripts annotated in the reference genome QQ74-V2. We compared accessions, day-length conditions, and developmental stages (Figure 1) (dgeFitGLM, edgeR). Second, we used novel transcripts to compare accessions, day-length conditions, and developmental stages, as done for the annotated transcripts. Later, those DEGs were used to conduct co-expression analyses, as described in the next section.

**Figure 1.**
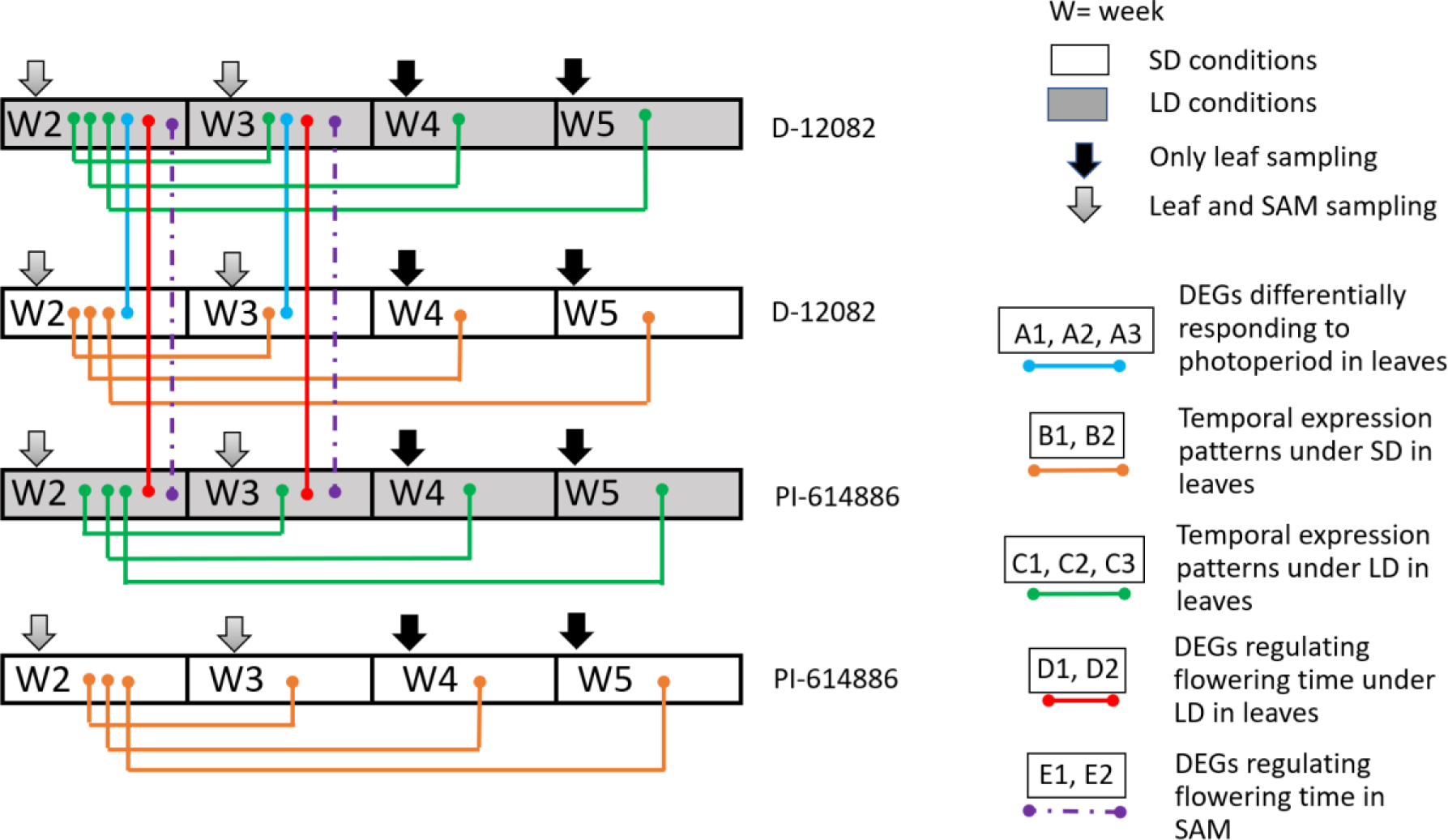
Schematic view of the experimental design, sampling points, and relevant comparisons between accessions, day length regimes, and developmental stages to identify putative candidate genes. Different comparisons are described using the identifiers in Table 1. Plants were grown in a growth chamber at 22°C under long-day (LD, 16 h light) and short-day conditions (SD, 8 h light). Samples were harvested weekly at ZT-9 (leaves) and ZT-9 to ZT-12 (SAM). W=week.

**Table 1.**
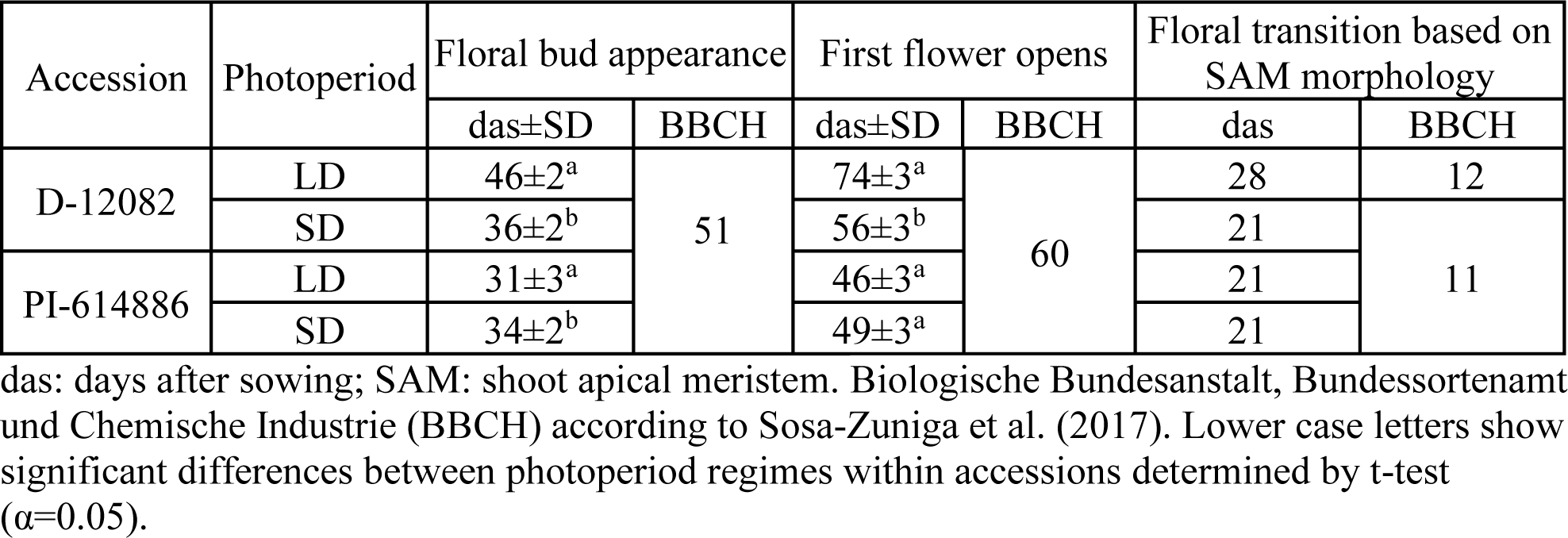
Macroscopic and microscopic observations on floral transition in two quinoa accessions. D-12082 is a short-day accession from Peru, and PI-614886 is a day-neutral accession from Chile. Plants of each accession were grown in a growth chamber at 22°C under long-day (LD, 16 h light) and short-day conditions (SD, 8 h light).

### Co-expression analysis

We used co-expression networks to associate novel transcripts to a putative biological process. For this purpose, we combined the DEGs between accessions, day-length conditions, and developmental stages obtained for the novel transcripts with the results from the annotated DEGs (dgeFitGLM, edgeR). Then, we used the R package BioNERO (Almeida-Silva & Venancio, 2022) to perform co-expression analyses in the data set containing annotated and novel DEGs. First, we calculated pairwise Pearson correlation coefficients between gene expression patterns. Then, we assessed the interaction between different genes to construct modules using the Weighted Gene Co-expression Network Analysis (WGCNA) algorithm (signed-hybrid network with default parameters) (scale-free power β=11 of threshold 0.8, module merging threshold=0.8; depth split=10) (Langfelder & Horvath, 2008). Later, we constructed co-expression networks for each of the modules. Then, we selected the top 10% of novel transcripts with the highest degree and a module membership (MM) > 0.7. A degree is defined as the sum of the connection weights of a gene to all other genes in the module, while MM is the correlation of a gene to its module eigengene (Almeida-Silva & Venancio, 2022). As a last step, we blasted the selected novel transcripts using the NCBI Basic Local Alignment Search Tool (BLAST) service to perform blastx (Taxid: 58023; E-value: 1.0E-3, number of Blast Hits: 3, word size: 6).

### Gene Ontology annotation

We carried out Gene Ontology (GO) annotation of the resulting DEGs for each of our analyses (annotated and novel DEGs). First, we used the NCBI BLAST service as described above (Taxid: 58023; E-value: 1.0E-3, number of Blast Hits: 3, word size: 6). We used the blastx results as input to Blast2GO (version 6.0) to map and annotate GO terms (annotation cut-off: 55, GO-weight: 5) (Götz *et al*., 2008). Later, we performed Enzyme Code annotation and Kyoto Encyclopedia of Genes and Genomes (KEGG) mapping with the Blast2GO tool (Kanehisa & Goto, 2000) using the GENES and PATHWAYS data bases (http://www.genome.ad.jp/kegg/). When the blastx results of a novel transcript indicated a similarity to a gene previously described in the literature as involved in photoperiod response or flowering time regulation, we investigated the domains of the translated protein *in silico* (https://www.ncbi.nlm.nih.gov/Structure/cdd/wrpsb.cgi).

### Real-time qPCR

DNase-treated RNA was reverse transcribed with the First Strand cDNA Synthesis Kit (Thermo Fisher Scientific Inc., Waltham, United States), according to the manufacturer’s instructions. We performed RT-qPCR by Bio-Rad CFX96 Real-Time System, which has a built-in Bio-Rad C1000 Thermal Cycler (Bio-Rad Laboratories GmbH, Munich, Germany). We used 18 μl of master mix and 2 μl of diluted (1:20) cDNA per reaction. The master mix composition was as follows: 10 μl of Platinum SYBR Green qPCR SuperMix-UDG with ROX (Invitrogen by Life Technologies GmbH, Darmstadt, Germany), 1 μl of forward primer, 1 μl of reverse primer, and 6 μl of ddH_2_O. The amplification conditions were as follows: 95°C for 3 min as initial denaturation and 40 cycles of 10 s at 95°C, 20 s at primer pair annealing temperature, and 30 s at 72°C. We amplified three technical replicates per cDNA sample and used water as a template for negative control reactions. Cq values were obtained by setting the baseline threshold to 100, and expression levels were calculated by the comparative ΔΔCt method (Livak & Schmittgen, 2001). Primers used for gene expression analysis are listed in Supplementary Table 2. *CqPTB* and *CqIDH-A,* identified in our previous study as suitable housekeeping genes for RT-qPCR analysis in quinoa, were used as reference genes (Maldonado-Taipe *et al*., 2021).

### Statistical analyses

Pearson’s R correlation was calculated using MS Excel between the expression values obtained by RNA-seq (dgeFitGLM data contrasts) and those obtained by RT-qPCR (ΔΔCt method). DTB and DTF significant differences between photoperiod regimes were determined by t-tests (α=0.05). The correlation between DTF and DTB was calculated and reported as Pearson correlation coefficient (r).

## Results

### Histological investigation of the quinoa shoot apical meristem

Considering the flowering time as “first flower opens”, the accession PI-614886 flowered much earlier than D-12082 under both SD (seven days earlier) and LD (18 days earlier) conditions, as expected from a previous study (Patiranage *et al*., 2021) (Table 1). However, both accessions bolted simultaneously under SD conditions, while PI-614886 bolted much earlier than D-12082 under LD. Nevertheless, the opening of the first flower was highly correlated with bolting time in both accessions under LD (r=0.57 and r=0.75) and SD (r=0.89 and r=0.47). Interestingly, D-12082 plants generally showed more vegetative growth (bigger leaves and more branches) under LD than under SD conditions.

We analyzed the apices under the microscope to determine when the SAM is turned into a floral meristem under LD and SD in both accessions. We defined the time of floral transition when the SAM exhibited a “dome shape” phenotype, which corresponds to the SAM morphology at the reproductive stage (apical dominance release) (Dun *et al*., 2006) (Figure 2). This “dome shape” was observed when the plants were at a relatively early stage of development, at BBCH11 (first pair of leaves visible) and BBCH12 (second pair of leaves visible) (Sosa-Zuniga *et al*., 2017) (Supplementary Figure 1). Under SD conditions, PI-614886 and D-12082 displayed floral transition three weeks after sowing whereas, under LD, PI-614886 and D-12082 exhibited floral transition at different time points, three and four weeks after sowing, respectively (Table 1).

**Figure 2.**
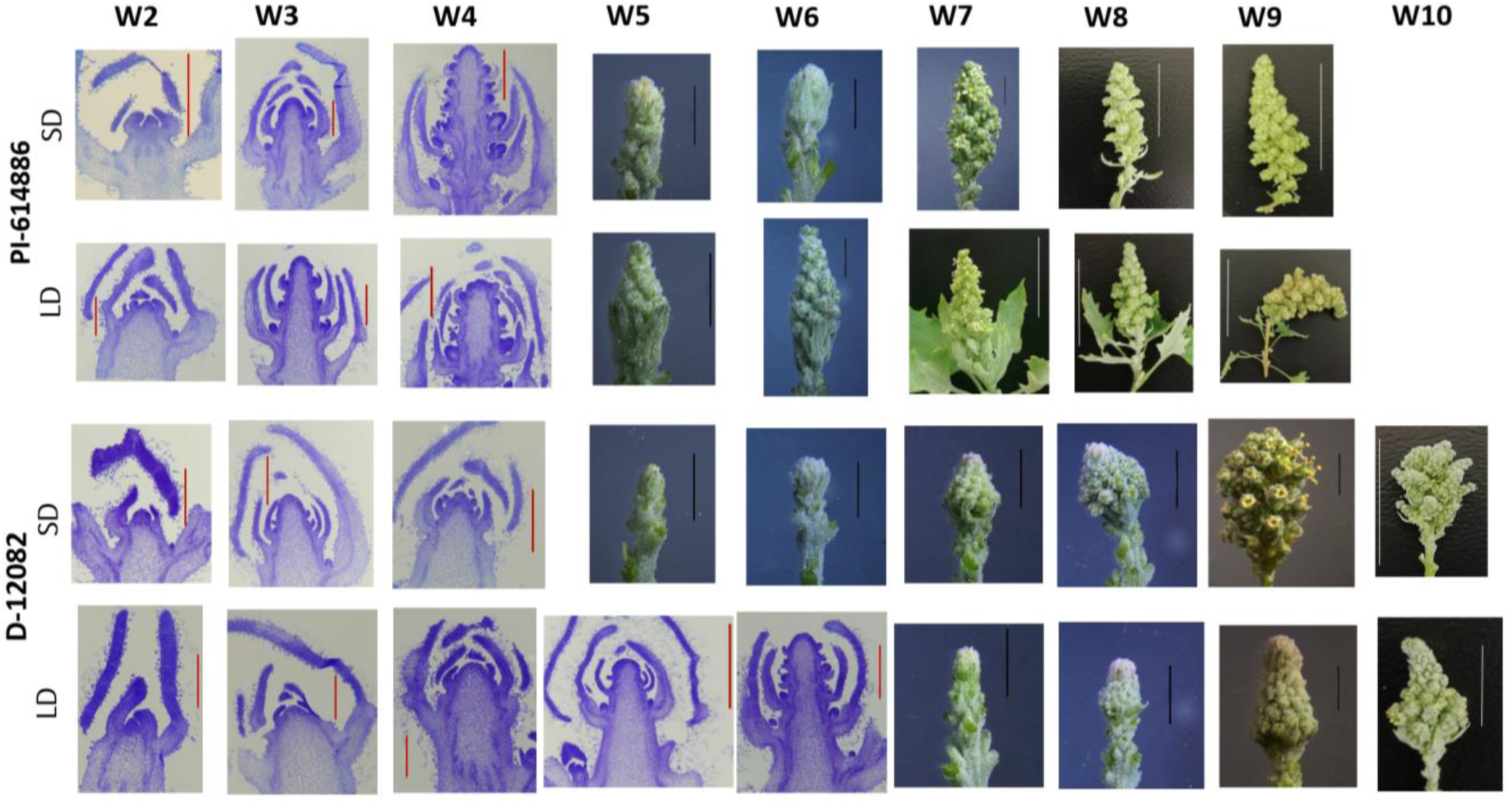
Shoot apical meristem (SAM) and inflorescence development from two quinoa genotypes. PI-614886 and D-12082 are day-neutral and short-day accessions, respectively. The SAM longitudinal sections were stained with toluidine blue. Plants were grown in a growth chamber at 22°C under long-day conditions (LD, 16 h light), and short-day conditions (SD, 8 h light). Photos were taken two to 10 weeks after sowing (W2-W10). A vegetative SAM was identified by an “edged” shape (e.g., W2 accession D-12082, LD), whereas a reproductive meristem was identified by a “dome shape” (e.g. W4 accession D-12082, LD). Scale bars: 500 µm (red), 5000 µm (black), 37.5 mm (white).

### Transcriptome sequencing

We sequenced 48 leaf and 24 SAM cDNA libraries from plants grown under SD and LD conditions. In total, 68.66 million reads per library were generated with an average read length of 150 bp, amounting to 20.59 Gbp per library (Supplementary Table 3). We discovered 132,965 gene models, of which, after excluding the unexpressed genes (those with about five read counts or more in a minimum number of samples), 85,634 gene models in leaf and 74,640 gene models in SAM were retained. A total of 71,230 genes were present in both SAM and leaf tissues. Of the retaining gene models, 26,782 were annotated in the reference genome QQ74-V2 (CoGe Genome ID: id60716), and 57,342 were novel transcripts.

We constructed MDS plots with the filtered and normalized leaf and SAM transcriptomes. Following our expectations, we observed that the samples of the same biological replicates corresponding to the same tissue/treatment clustered together (Supplementary Figure 2). Dimension 1 explained the most variation (19.0%) separating the SAM samples from the leaf samples. Dimension 2 separated the PI-618886 and D-12082 samples and explained 13.0% of the variation-(Supplementary Figure 2).

### Identification of differentially expressed genes with a putative function as flowering time regulators

First, we analyzed the transcripts annotated in the reference genome QQ74-V2. We aimed to identify gene expression profiles that could correlate to the meristem morphological changes. We focused on BBCH11 and BBCH12 (W2 and W3, respectively) developmental stages because our study of the SAM at the histological level showed that floral transition occurred at those stages. We analyzed our transcriptome data in four steps. First, we looked for genes differentially responding to photoperiod to regulate flowering time in leaves. Second, we looked for genes regulating flowering time under LD conditions in leaves. Third, we searched for genes regulating the time to flower under SD conditions in leaves. Finally, we screened for the DEGs in SAM to identify flowering time candidate genes in this tissue.

### DEGs responding to photoperiod in leaves

To discover genes differentially responding to photoperiod, we investigated the DEGs between SD and LD conditions at W2 and W3 in D-12082 (solid blue lines in Figure 1) because the morphological changes at the SAM indicated that transition to a floral meristem in this accession occurs in a photoperiod-dependent manner (Supplementary Figure 3A). We selected the genes for which the expression changes between SD and LD in W2 were significantly different compared to those in W3, given the substantial morphological differences of SAMs between W2 and W3 (accession D-12082). For this purpose, we used the criteria −0.25≥(log_2_ FC at W2 – log_2_ FC at W3) ≥0.25 (Table 2, Supplementary Figure 3B).

**Table 2.**
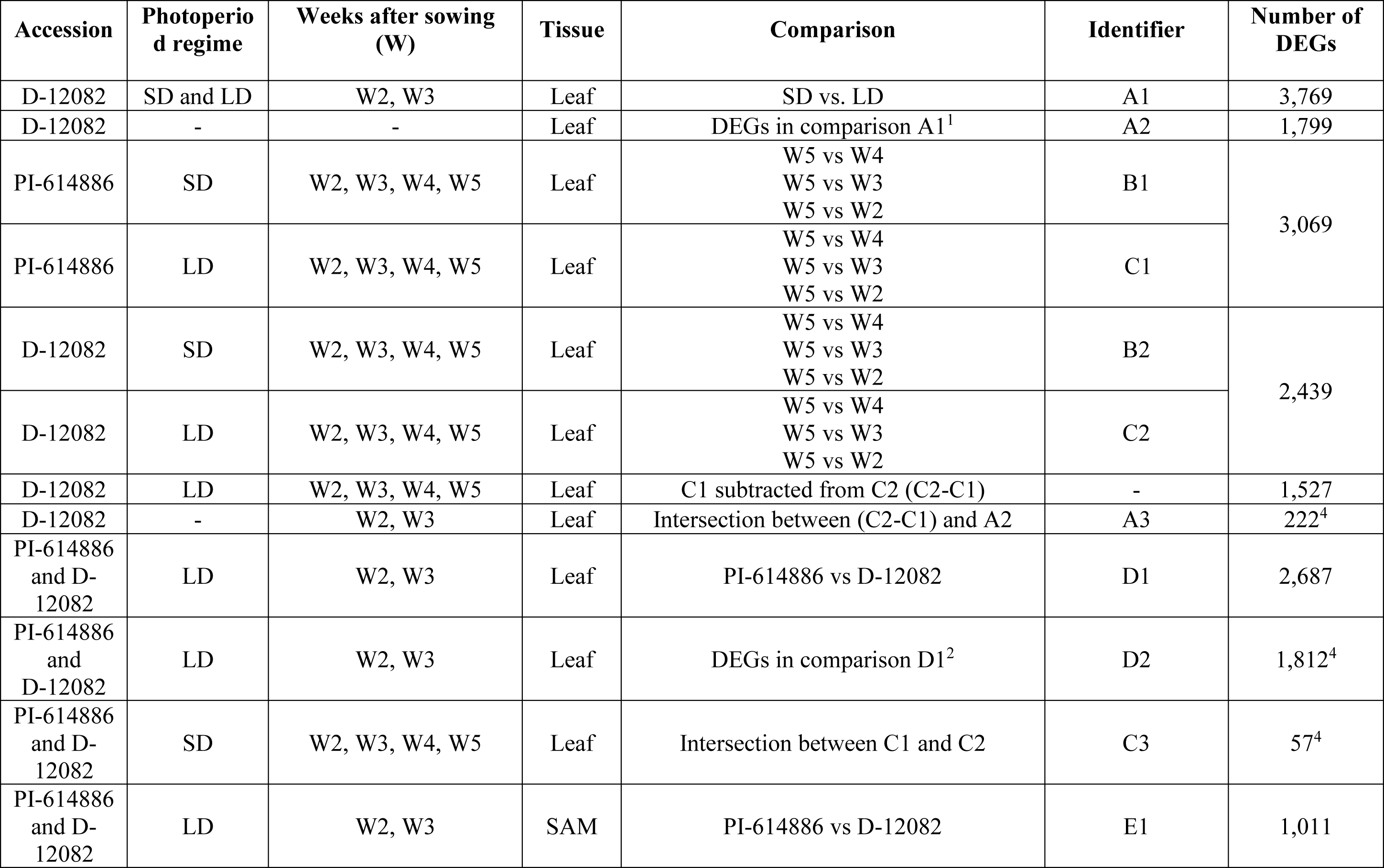
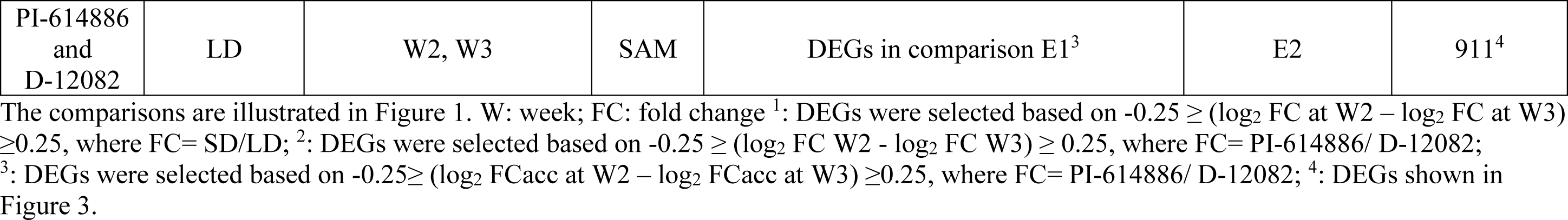
Summary of differentially expressed genes (DEGs) found by transcriptome sequencing. RNA was taken from plants grown in a growth chamber under short- and long-day conditions.

Subsequently, we investigated the temporal expression pattern of the selected DEGs throughout the plant’s development in both accessions under LD and SD (green solid lines and orange solid lines in Figure 1, respectively) (Supplementary Figure 3C and Supplementary Figure 3D). Based on the morphological changes of the SAM, we expected that flowering time genes differentially responding to photoperiod show different temporal expression patterns between SD and LD in the photoperiod-sensitive accession, D-12082, but not in the day-neutral accession, PI-618886. Therefore, we subtracted the DEGs in the day-neutral accession from those in the short-day accession (Supplementary Figure 3E). Finally, we selected those DEGs that resulted from the subtraction and were also differentially expressed between LD and SD in D-12082 (solid blue lines in Figure 1; Supplementary Figure 3B). As a result, we selected 222 genes differentially responding to photoperiod, which putatively control flowering time in leaves (Table 2, Figure 3). Among the resulting DEGs and based on a literature survey, we found several genes with a known function in photoperiodic regulation of flowering time in other species like *TREHALOSE-PHOSPHATASE/SYNTHASE 9* (*CqTPS9*), *PHYTOCHROMEB* (*CqPHYB*), *ZEITLUPE* (*CqZTL*) and *BLUE-LIGHT INHIBITOR OF CRYPTOCHROMES* (*CqBIC1*) (Figure 3 and Supplementary Table 4) (Blázquez & Weigel, 1999; Más *et al*., 2003; Wang *et al*., 2016; Tian *et al*., 2021).

**Figure 3.**
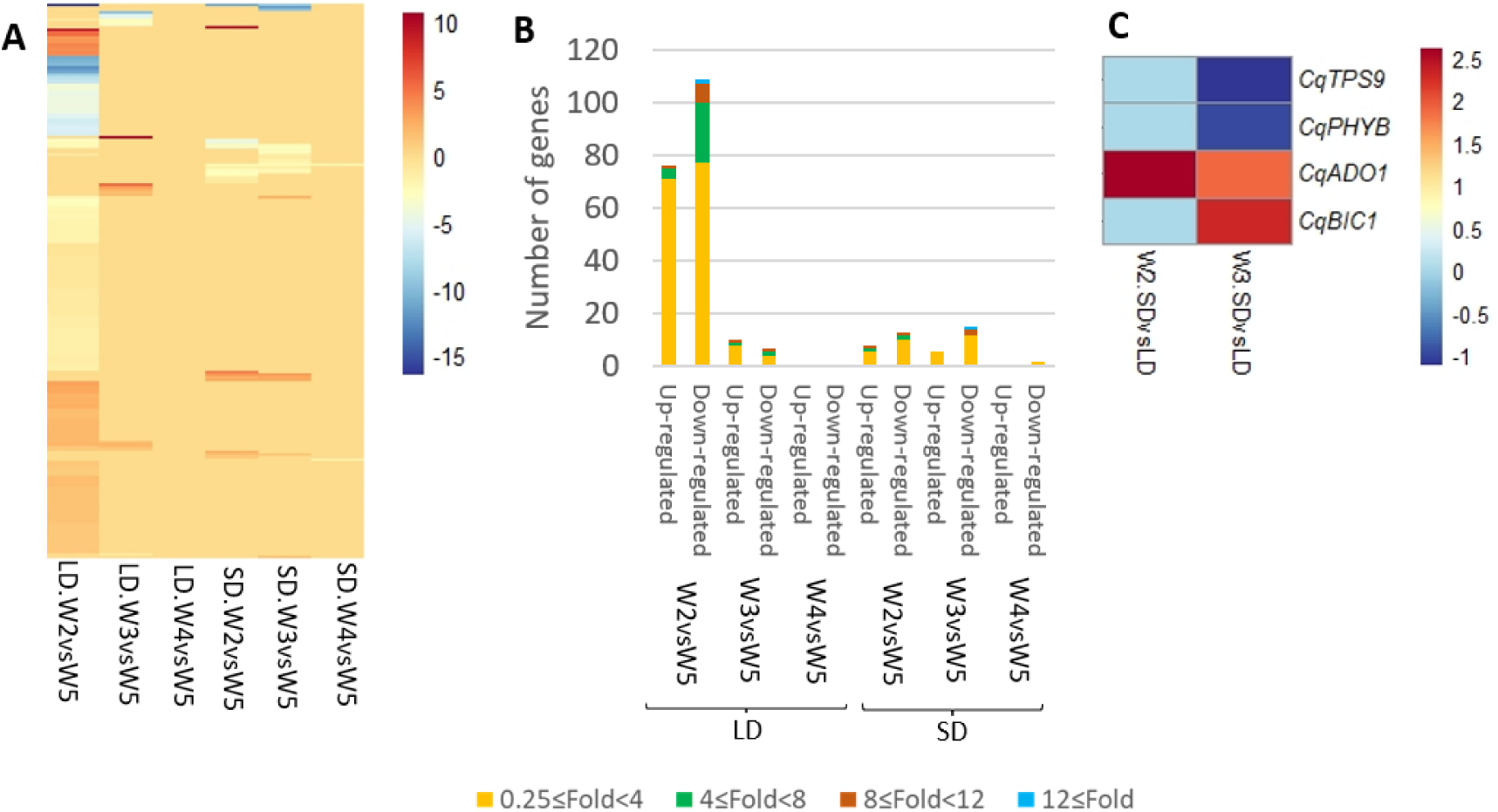
DEGs in D-12082 that differentially respond to photoperiod to likely regulate flowering time (identifier A3 in Table 1). (A) Heatmap, (B) number of up- and down-regulated genes, and (C) selected genes with a known flowering time-related function in other species. Plants were grown under long and short day (LD, 16 h light) conditions (SD, 8 h light). RNA was isolated from leaves. LD.W2vsW5 corresponds, for instance, to log_2_ (normalized reads in W2/ normalized reads in W5). The heatmap was constructed with the pheatmap R package. W=week.

### DEGs regulating flowering time under LD in leaves

As a next step, we searched for genes expressed in the leaves, which putatively regulate flowering time under LD conditions (red solid lines in Figure 1) since we considered the possibility of different flowering time regulators under SD and LD conditions. These genes could be downstream targets of the DEGs differentially responding to photoperiod. We assumed that the main flowering time integrators are the same for the day-neutral and the short-day accessions, implying that a gene integrating flowering signals would not be accession-dependent. Accordingly, we investigated the DEGs between PI-614886 and D-12082 under LD at W2 and W3 (red solid lines in Figure 1, Supplementary Figure 4). From the obtained DEGs, we kept genes with −0.25 ≥ (log_2_ FC W2 - log_2_ FC W3) ≥ 0.25, whose expression pattern was significantly changed between W2 and W3. The obtained down- and up-regulated genes at long-day conditions at week 2 (referred to as LD.W2 in Figure 3) could serve as putative flowering repressors and promoters, respectively (Figure 4, Table 2). Among the obtained 1,812 DEGs, we found several genes previously reported as flowering time regulators in quinoa and other species (Supplementary Table 4), for example *FLOWERING LOCUS T 1* (*CqFT1A*)*, FLOWERING LOCUS T 2* (*CqFT2B*)*, HEADING DATE 3A* (*CqHD3AB*)*, CONSTANS-LIKE 16* (*CqCOL16*)*, CONSTANS-LIKE 4* (*CqCOL4*)*, FRIGIDA INTERACTING PROTEIN (CqFIP1), FRIGIDA* (*CqFRL4A*)*, EARLY FLOWERING 5* (*CqELF5*) (Suarez-Lopez *et al*., 2001; Kim *et al*., 2008; Komiya *et al*., 2008; Pin *et al*., 2010; Choi *et al*., 2011; Patiranage *et al*., 2021).

**Figure 4.**
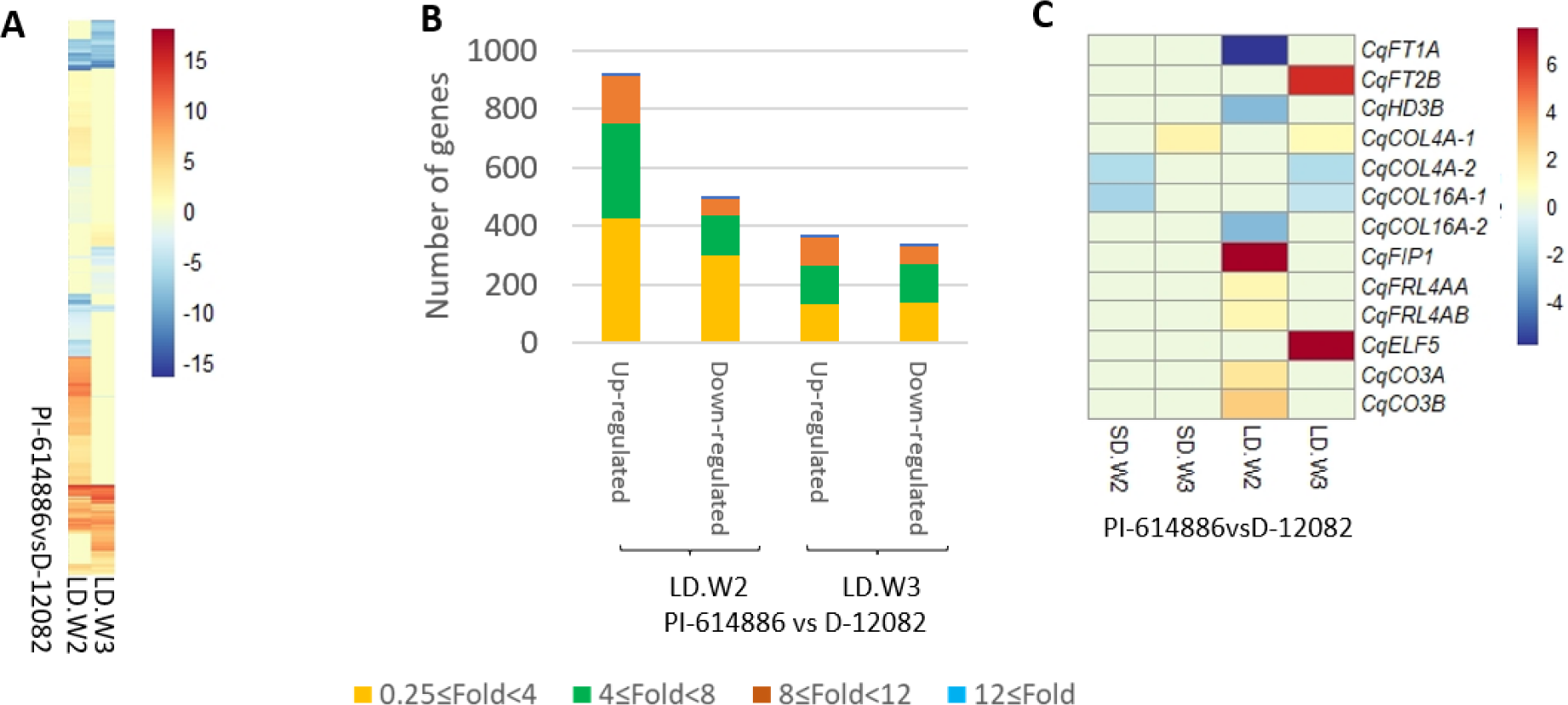
DEGs between PI-614886 and D-12082 at W2 and W3 under long day (LD) conditions putatively control flowering under LD (identifier D2 in Table 1). (A) Heatmap, (B) number of up- and down-regulated genes, and (C) selected genes with a known flowering time-related function in other species. Plants from the accessions PI-614886 and D-12082 were grown under long-day conditions (LD, 16 h light), and short-day conditions (SD, 8 h light). RNA was isolated from leaves. LD.W2.PI-614886vsD-12082 corresponds, for instance, to log_2_ (normalized reads in PI-614886/ normalized reads in D-12082). The heatmap was constructed with the pheatmap R package. W=week.

### DEGs regulating flowering time under SD in leaves

In the next step, we searched for genes expressed in leaves that likely regulate flowering time under SD conditions. Since morphological changes at the SAM indicate that both accessions transit to flowering at the same time under SD, we searched for the DEGs in the day-neutral accession with similar temporal expression patterns to those of the short-day accession (orange solid lines in Figure 1). Among the 57 resulting genes, we found *SUPPRESSOR OF CONSTANS OVEREXPRESSION 1* (*CqSOC1*) (Figure 5 and Supplementary Table 4), which integrates multiple flowering signals derived from the age-dependent and gibberellin pathways in *Arabidopsis* and other species (Lee & Lee, 2010).

**Figure 5.**
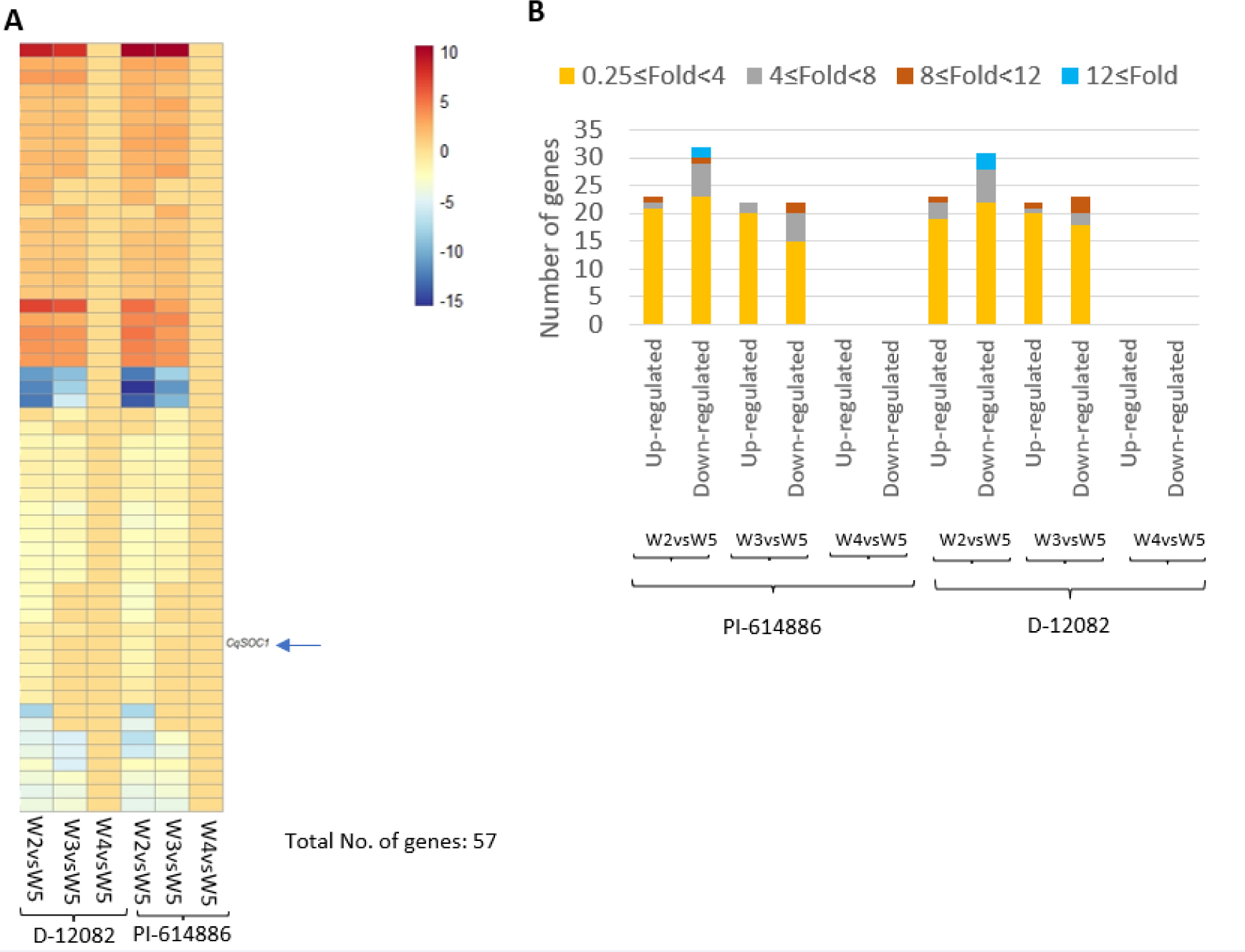
DEGs with similar temporal expression patterns in PI-614886 under short-days (SD) putatively control flowering under SD (identifier C3 in Table 1). (A) Heatmap and (B) number of up- and down-regulated genes. An arrow shows the expression pattern of *SOC1*. Plants were grown under long-day conditions and short-day conditions (SD, 8 h light). RNA was isolated from leaves. LD.W2vsW5 corresponds, for instance, to log_2_ (normalized reads in W2/ normalized reads in W5). The heatmap was constructed with the pheatmap R package. W=week.

### DEGs regulating flowering time in SAM

Lastly, we searched for genes regulating flowering time in the SAM. Here we investigated the DEGs between accessions under LD at W2 and W3 when differences in time of floral transition occurred at the SAM (purple dashed lines in Figure 1; Supplementary Figure 5). Given the morphological differences observed in the sections of the SAM, the expression fluctuations between the accessions (log_2_ FC_acc_) must be different between W2 and W3 (−0.25≥ (log_2_ FC_acc_ at W2 – log_2_ FC_acc_ at W3) ≥0.25)). In total, 911 DEGs showed differential expression between the accessions under LD conditions but not under SD, where both accessions flowered simultaneously (Figure 6 and Table 2). Interestingly, among these genes, which putatively regulate flowering time in quinoa in the SAM, we found homologs of known flowering time regulators in other species (Supplementary Table 4). A quinoa homolog of *AGAMOUS-LIKE 15* (*CqAGL15*), a flowering repressor in Arabidopsis (Adamczyk *et al*., 2007), was down-regulated at W2 under LD conditions only in PI-614886 but not in D-12082. A quinoa homolog of *APETALA1* (*CqAP1*), a flowering regulator in Arabidopsis, soybean, wheat, and other species (Murai *et al*., 2003; Chen, L *et al*., 2020), was up-regulated in PI-614886 at W3 under LD conditions. Moreover, a homolog of *FT INTERACTING PROTEIN 3* (*CqFTIP3*), required to maintain floral meristems in Arabidopsis (Liu *et al*., 2018), was down-regulated at W3 in PI-614886 compared to D-12082.

**Figure 6.**
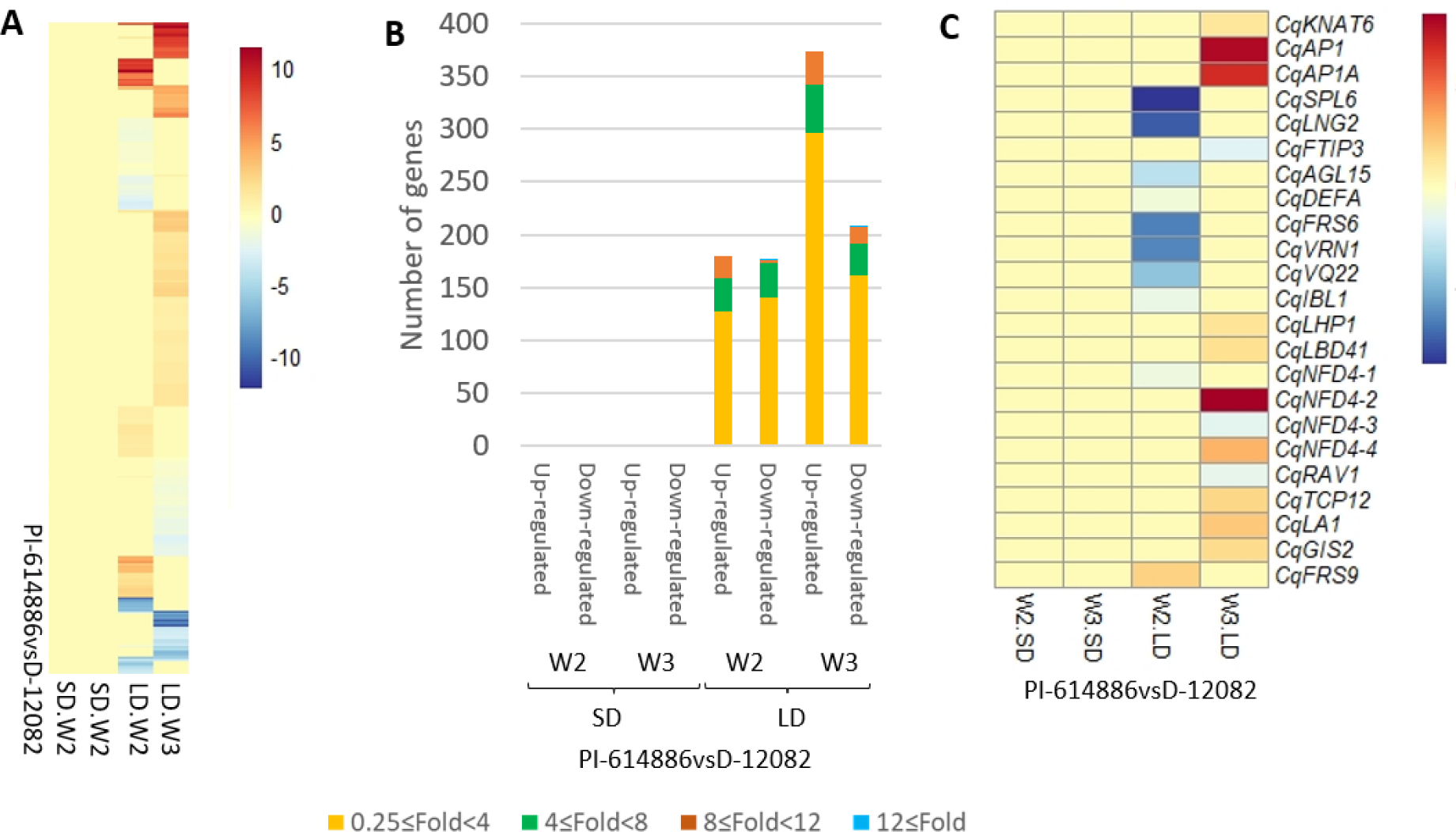
DEGs between PI-614886 and D-12082 at W2 and W3 are likely to regulate the time to flower at the shoot apical meristem (SAM) (identifier E2 in Table 1). (A) Heatmap, (B) number of up- and down-regulated genes, and (C) selected genes with a known flowering time-related function in other species. Plants from the accessions PI-614886 and D-12082 were grown under long-day conditions (LD, 16 h light) and short-day conditions (SD, 8 h light). RNA was isolated from SAM tissue. LD.W2.PI-614886vsD-12082 corresponds, for instance, to log_2_ (normalized reads in PI-614886/ normalized reads in D-12082). The heatmap was constructed with the pheatmap R package. W=week.

### Gene ontology of differentially expressed genes

We analyzed the putative function of all DEGs identified in this study. The 222 DEGs which differentially respond to photoperiod were mainly allocated to “Regulation of gene expression” and “Transcription, DNA-templated” (GO terms: 0010468 and 0006351) under the “Biological Process” category (Supplementary Table 5). Furthermore, 56 genes (25.22%) were found in the KEGG database and were allocated to38 signaling pathways: glycine, serine, and threonine metabolism (7.14%), glycolysis/gluconeogenesis (5.36%), among others (Supplementary Table 6). The percentage of DEGs found in the KEGG database varied from 5.96 to 28.81% when analyzing the DEGs that likely regulate flowering time in the SAM and, under LD and SD conditions, in the leaves (Supplementary Tables 7 to 12).

### Co-expression analysis of the DEGs

We performed a co-expression analysis with the novel transcripts and genes annotated in the reference genome QQ74-V2. We hypothesized that genes within the same co-expression cluster might have similar functions. First, we performed a differential expression analysis with the putative novel transcripts as described for the annotated genes (steps described in Table 2 were also carried out for novel transcripts). Second, we grouped the resulting transcripts from the differential expression analysis as follows: novel (89) and annotated (222) DEGs differentially responding to photoperiod (data set 1); novel (1,777) and annotated (1,812) DEGs that might regulate flowering time under LD conditions (data set 2), novel (127) and annotated (57) DEGs that might regulate flowering time under SD conditions (data set 3), and novel (6,229) and annotated (911) DEGs that likely regulate flowering time in the SAM (data set 4). Third, we performed separate co-expression analyses with these four data sets. As an outcome, we identified six co-expressed modules with data set 1 (Figure 7). The red module contained the highest number of DEGs (88 DEGs) (Supplementary Figure 6). Moreover, we identified 20, three, and 12 co-expressed modules with data sets 2, 3, and 4, respectively (Figure 7). The number of DEGs harbored by a module ranged between 72 and 1,957 (Supplementary Figure 6). Interestingly, the grey module of data set 2 contained 884 novel DEGs with *CqFT1A, CqFT2B, CqCOL16,* and *CqELF5*. Furthermore, in data set 3, the grey module clustered 56 novel DEGs with *CqSOC1*.

**Figure 7.**
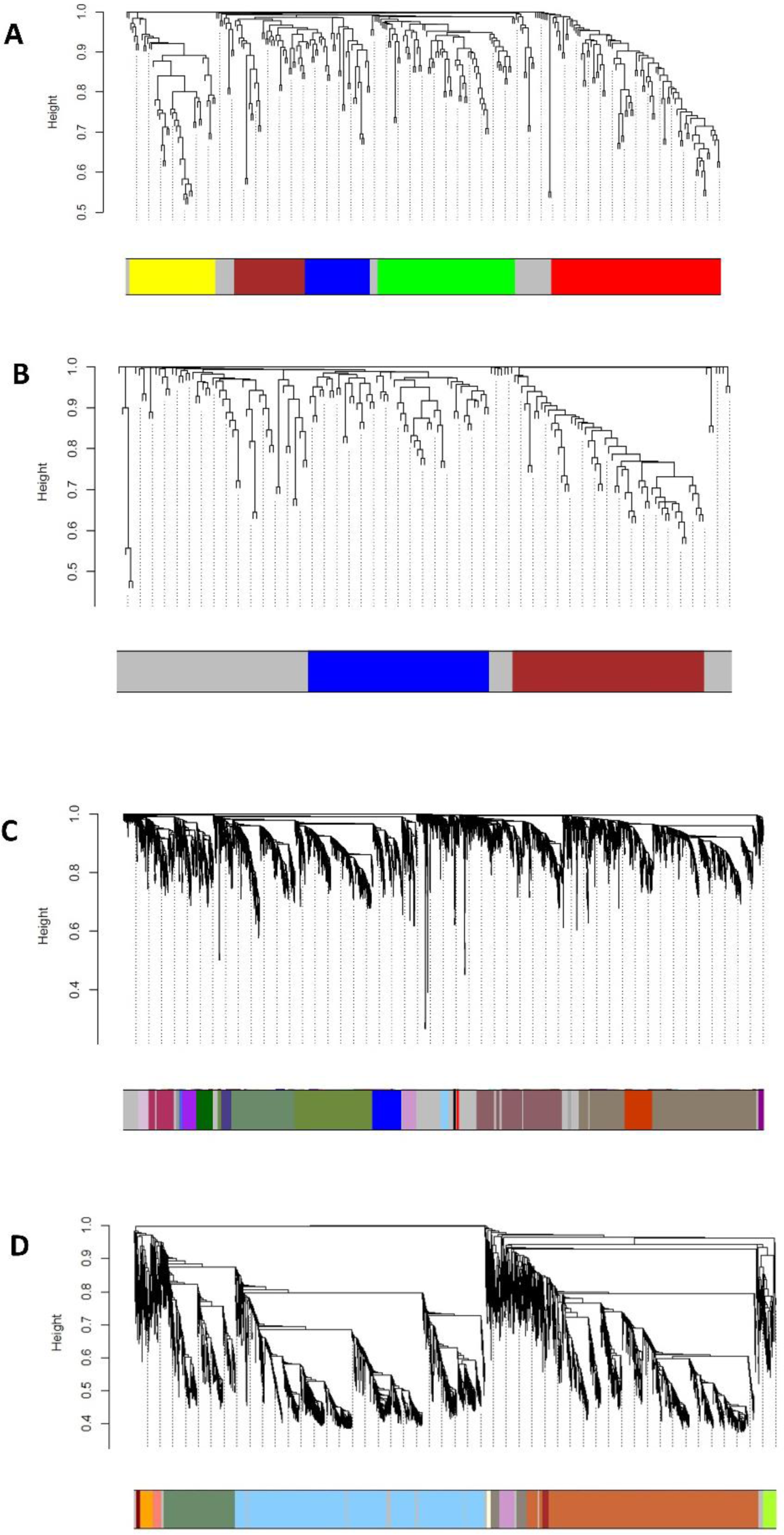
Co-expression analysis of annotated and novel (non-annotated) DEGs in leaves. (A) Differentially expressed genes (DEGs) between short-day (SD) and long-day (LD) conditions in D-12082, which differentially respond to photoperiod to likely regulate flowering time (identifier A3 in Table 1 and Figure 1). (B) DEGs between PI-614886 and D-12082 at W2 and W3 under LD are putatively controlling flowering under LD (identifier D2 in Table 1 and Figure 1). (C) DEGs with similar temporal expression patterns between PI-614886 and D-12082 likely control the time to flower under SD (identifier C3 in Table 1 and Figure 1). (D) DEGs between accessions with a putative function of flowering time regulation in the SAM (identifier E2 in Table 1 and Figure 1). Hierarchical clustering dendrograms of the DEGs are shown at the top. Every cluster (module) is shown by a different color at the bottom of the dendrograms. The y-axis (Height) displays the distance between clusters.

### Weighted Gene Co-expression Network Analysis

As a next step, we constructed co-expression networks of each module (Supplementary Figure 7 and Supplementary Figure 8). We selected 550 novel leaf and 616 SAM transcripts for further examination, representing the top 10% of transcripts in each module with the highest degree and MM > 0.7. We blasted and GO-annotated these novel transcripts to further characterize their putative function (Supplementary Table 13 and Supplementary Table 14). As an outcome, by BLAST analysis, 84.18% of the novel transcripts in leaves depicted sequence similarity to one or more gene orthologs in other species, out of which 117 (25.27%) coded for uncharacterized proteins. In the case of the SAM tissue, we found that 75.16% of the novel transcripts had sequence similarity to one or more gene orthologs in other species, out of which 177 (38.22%) coded for uncharacterized proteins (Supplementary Table 13). In total, 148 selected transcripts were annotated to be an isoform of a known gene, and 61 had sequence similarity to genes previously described in the literature as flowering time or photoperiod regulators (Supplementary Table 15). For instance, we found sequences with similarity to *RICESLEEPER*, whose homologs were associated with days to flowering in quinoa (Maldonado-Taipe *et al*., 2022) and to *APETALA 2-like* (*AP2*), a flowering time regulator in several species (Debernardi *et al*., 2020; Shim *et al*., 2022). We also found sequences with similarity to the Arabidopsis genes *FLOWERING LOCUS M* (*FLM*), whose splicing variants are known to repress flowering time (Scortecci *et al*., 2001; Lutz *et al*., 2015), and to ZTL, which modulates the circadian rhythm (Más *et al*., 2003).

### Validation of the RNA-seq results by RT-qPCR

We validated our transcriptome results by performing RT-qPCR analyses with paralogs of *CqTPS9*, *CqZTL*, *CqFIP1*, *CqELF5, CqSOC1, CqFT2B,* and *CqHD3AB* (Supplementary Table 4). These genes were randomly selected from the list of candidate genes because they are homologs of known flowering time regulators. Then, we calculated the correlation coefficient between the differential expression values from RNA-seq and the ΔΔCt values from RT-qPCR. The differential expression analyzed by RT-qPCR closely matched the RNA-seq results (r=0.82) (Supplementary Figure 9). Moreover, the expression patterns obtained by RT-qPCR matched our expectations based on our transcriptome data dissection (Figure 8). For instance, *CqZTL* relative expression is higher in D-12082 than in PI-614886 under LD at W3. Thus, *CqZTL* expression is higher in the accession that differentially responds to photoperiod at the stage when flowering commences in LD, based on our histological analysis. These results align with a gene that responds differently to photoperiod to control flowering time. Moreover, the relative expression of *CqZTL* in PI-614886 remains relatively unchanged under both day-length conditions, as expected from PI-614886, a day-neutral accession according to our histological analysis. *CqFT2B* provides another example. This gene’s expression was much higher in PI-614886 than in D-12082 under LD at W3, matching our RNA-seq observations (Figure 4). Interestingly, the relative expression of *CqFT2B* was higher towards W3 under SD in both accessions, aligning with the floral transition time as detected in our histological analysis (BBCH11).

**Figure 8.**
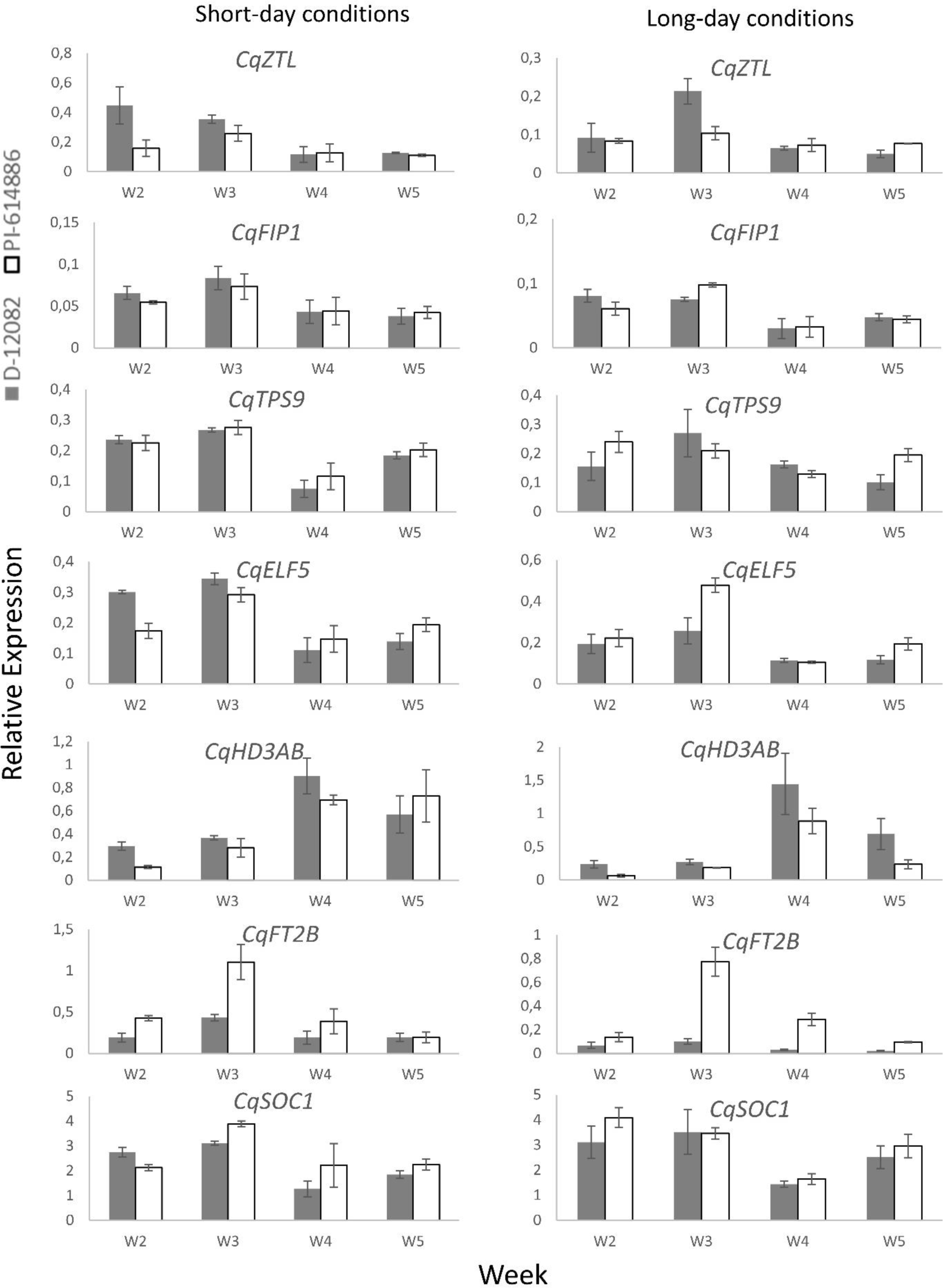
Leaf expression profiles of seven genes at different stages of development were obtained by RT-qPCR. Plants from accessions PI-614886 and D-12082 were grown in a growth chamber at 22°C under short-day (SD, 8 h light) and long-day conditions (LD, 16 h light). Three biological replicates and three technical replicates of each biological replicate were used. Error bars: ±*SEM*; data were normalized against the geometric mean of *CqPTB* and *CqIDH-A*. W=week.

## Discussion

We present a comprehensive study about the floral transition in quinoa encompassing morphological, phenological, and gene expression data. Two accessions with contrasting day-length responses were selected from previous experiments. By histological analysis, we determined when the SAM became a floral meristem. We identified 222 genes differentially responding to photoperiod and 1,812 and 57 genes putatively regulating flowering time under LD and SD conditions, respectively. We also identified 911 genes that putatively regulate flowering time in the SAM.

The histological results placed the floral transition time in quinoa much earlier than the bolting time. We conclude that genes triggering floral transition should be already active at the early stages of quinoa development when the second pair of leaves appears (BBCH12). This agrees with reports in *Chenopodium rubrum,* a species closely related to quinoa, where *CrFTL* genes respond to SD conditions to flower as early as five days after sowing (Cháb *et al*., 2008). Although floral transition starts early in quinoa, the onset of flowering (when the first flower opens) was recorded several weeks after the floral transition occurred at the SAM, demonstrating that the time to floral transition alone is not the sole predictor of days to flowering. In this case, the time from floral transition until the first flower opens is affected by several environmental factors, as demonstrated in several crops (e.g., tomato, rice, wheat), where nutrient supply, day length, light intensity, light quality, and ambient temperature, as well as endogenous signals transmitted by plant hormones play an important role (Camejo *et al*., 2005; Bäurle & Dean, 2006; Hu *et al*., 2015).

We searched for genes likely involved in photoperiod response and flowering time regulation based on their differential expression between accessions, day-length conditions, and developmental stages. There are constitutively expressed flowering time regulators, such as the *FTL* homolog in Chrysanthemum (Higuchi & Hisamatsu, 2015). However, the reason for the experimental design in our study was that many flowering time genes differentially respond to photoperiod, and their expression is correlated with the morphological changes in the SAM (Putterill & Varkonyi-Gasic, 2016). Other possible experimental designs include studying diurnal expression patterns and/or switching between photoperiod regimes. Such experimental settings allowed Wu *et al*. (2023) to identify *CO-like* transcription factors sensitive to the quinoa switch from LD to SD. Moreover, introducing a night break and studying lncRNA instead of mRNA in quinoa, as done by Wu *et al*. (2021), would represent another experimental design that would unveil novel candidate genes putatively involved in photoperiod response.

None of the genes found to be down- or up-regulated after NB by Wu *et al*. (2021) (*LHY, ELF3*, *HY5*, *PHYA*, and *CRY1*) were differentially regulated in our study. However, *CqFT2B* and *CqZTL*, identified by Wu *et al*. (2023) in a transcriptome study of diurnally collected samples under SD and LD, were also found in our study, which supports their role as flowering time regulators. Moreover, Patiranage *et al*. (2021) provided additional evidence that *CqFT2B*’s temporal and diurnal expression in early- and late-flowering quinoa accessions underpins its function as a flowering time regulator. Regarding the *CqFT1A* paralog, it has been reported that its expression pattern is not correlated with flowering time (Patiranage *et al*., 2021; Wu *et al*., 2023). However, we found that *CqFT1A* is likely to act as a flowering time repressor under LD. The different outcome of our study compared to previous studies might be due to the timing of the sampling, which was at least one week earlier than in other studies. This earlier sampling was shown to be beneficial, considering the very early transition of vegetative to reproductive stage in quinoa, identified in this study. Furthermore, Patiranage et al., 2021 found that *CqFT1A* haplotypes in a core collection of 276 quinoa accessions were correlated with flowering time. Under LD conditions, the *CqFT1Aa* haplotype was associated with early flowering, while the *CqFT1Ab* haplotype was correlated with late flowering (Patiranage *et al*., 2021). Interestingly, the PI-614886 and D-12082 accessions used in our study had the *CqFT1Aa* and *CqFT1Ab* haplotypes, respectively. Accordingly, we observed a down-regulation of *CqFT1A* at W2 in PI-614886, which flowered earlier than D-12082 under LD, as expected from a floral repressor gene. Noteworthily, the putative function of *CqFT1A* under LD conditions corresponds to *BvFT1* from sugar beet, another member of the *Amaranthaceae* family, where it inhibits flowering under LD (Pin *et al*., 2010).

The number of differentially expressed genes depends on the experimental design and plant material used, so thousands to tens of thousands of genes will likely change their expression in response to different environments (Sowiński *et al*., 2020). The number of genes in our study roughly agrees with those in other transcriptome analyses. For instance, comparing four developmental stages of *C. ficifolium* under LD and SD conditions revealed 6,096 differentially expressed genes (DEGs) (Gutierrez-Larruscain *et al*., 2022). Moreover, a transcriptome study in quinoa reported 1,817 and 8,870 genes possibly associated with floral induction under LD and SD, respectively (Gutierrez-Larruscain *et al*., 2022).

Positioning differentially expressed genes within flowering time QTL provides compelling evidence for their function as flowering time regulators. Twelve candidate genes from our study had been localized within a major flowering time QTL (Maldonado-Taipe et al., 2022) (Supplementary Table 16). Among these genes, we found a homolog of the Arabidopsis *VQ22* gene, encoding a VQ protein, which plays a vital role in plant growth, development, and response to environmental conditions, most likely by acting as cofactors of WRKY transcription factors. Moreover, Arabidopsis *vq22* mutants showed stunted growth (Cheng et al., 2012).

We hypothesize that most DEGs are involved in photoperiod-response and flowering time regulation in quinoa. There is increasing evidence that most phenotypes, previously thought to be simply inherited, are controlled by many genes, although with minor effects. The omnigenic paradigm even proposes that any trait is influenced by all genes expressed at a certain time point or developmental stage (Boyle *et al*., 2017; Tautz *et al*., 2020; Mathieson, 2021). Examples for complex traits (e.g., floral development, organ size) in maize, sunflower, and chili pepper (Díaz-Valenzuela *et al*., 2023) have been reported. In a large field study with maize under different environments, 39 and 36 QTL with minor effects were found for days to silking and days to anthesis, respectively (Buckler *et al*., 2009). This study showed that differences in flowering time among 5,000 inbred lines were not determined by a few genes with large effects but by the minor cumulative effects of numerous quantitative trait loci. Furthermore, according to the recently proposed omnigenic model, genes can be classified as core or peripheral in their association with a specific trait. Therefore, a DEG might not belong to flowering-related pathways but play an indirect role, e.g., in nutrient uptake or stress response (Cho *et al*., 2017). Accordingly, we found around a hundred DEGs predicted to function as “Transcription regulator activators” and “Regulators of gene expression”, which are commonly predicted functions for flowering time regulators (Song *et al*., 2015). However, several other functional categories, not expected to be related to flowering time, were also found, such as “endopeptidase activity”, “aspartic-type peptidase activity”, and “Udp-glycosyltransferase activity”. Therefore, our results provide further evidence of the omnigenic regulation of flowering time.

Regarding the omnigenic nature of flowering time control, transcriptomics is a powerful tool to identify paralogs that show transcriptional profiles similar to those of a flowering time gene described for other species. Due to quinoa’s polyploid nature, we expect neo- or sub-functionalization and gene silencing as in the case of the *PEBP* (*phosphatidyl ethanolamine-binding proteins*) gene family. Out of 24 sequence homologs, only five had been identified as putative *FT* orthologues (Jarvis *et al*., 2017; Patiranage *et al*., 2021). Only the leaf transcripts of *CqFT2B* and *CqFT1A* were detected in our study. Furthermore, only one *HD3A*, *SOC1,* and *AGL* paralog was identified as a putative flowering time gene based on its expression pattern. Furthermore, our study showed that out of 18 *CONSTANS-like* genes, only *COL16-A1*, *COL16-A2*, *COL4A,* and *COL4B* are putative flowering time candidate genes because they showed differential expression between the studied accessions at the time of floral transition (Supplementary Table 17 and Supplementary Figure 10).

Interestingly, we found genes with high sequence similarity to vernalization-responsive genes from other species, including *EARLY FLOWERING 5* (*ELF5*), *FRIGIDA INTERACTING PROTEIN (CqFIP1),* and *FRIGIDA-like* (*CqFRL4A*) in the leaf transcriptome and *CqVRN1* in the SAM tissue. These genes are known as vernalization-related genes in Arabidopsis, rapeseed, or wheat (Johanson *et al*., 2000; Yan *et al*., 2003; Noh *et al*., 2004; Wang *et al*., 2011). Since quinoa does not require vernalization to flower, it is tempting to speculate that these genes possibly underwent neo-functionalization. Hence, these genes might have adapted their function during evolution distinctly from other *Amaranthaceae* species, such as sugar beet, which requires vernalization to flower under LD. This crop provided a striking example of neo-functionalization as an *FT* homolog was turned into a floral repressor (Pin *et al*., 2010).

We found a surprisingly high number of novel transcripts (57,342), including algorithmic artifacts, splice variants, and genes specifically transcribed under our experimental conditions. Similar results were reported in a recent study by Zheng *et al*. (2022), where >50,000 novel transcripts were discovered under cold stress, including splicing variants and 5′ or 3′ extensions or truncations. More than 6,000 genes had not been annotated in the reference genome. It is important to note that one line used in our study differs from the line used for establishing the reference genome. Therefore, some novel transcripts might be line-specific splicing variants. Subsequently, with the co-expression analysis followed by blast and protein domain studies, we could identify promising novel candidate genes based on their association with the annotated candidate genes.

This study provides new insights into floral transition in quinoa by combining morphological and gene expression data. Differentially expressed genes located within the confidence interval of previously identified flowering time QTL are the most promising candidate genes for floral transition in quinoa. As a next step, the candidate genes should be functionally characterized to confirm their role as flowering time regulators in quinoa. In this scenario, overexpression and knockout of the candidate genes would provide the ultimate evidence of their function in quinoa. Currently, there is no reliable protocol for quinoa transformation. However, a recent virus-mediated transient expression (VIGS) protocol could enable functional studies in this crop (Ogata *et al*., 2021). Our study can also broaden the genetic variation utilized in quinoa breeding programs by identifying the beneficial haplotypes of the putative candidate genes and their integration into breeding programs through crosses to develop cultivars adapted to diverse environmental conditions.

## Data Availability Statement

The raw sequencing data of the 72 *C. quinoa* samples sequenced in this study are available on EBI-ENA under the study number PRJEB64679.

## Acknowledgments

We thank Bettina Rohardt, Federico Barbier, Monika Bruisch, and Florence Muraya for their technical assistance. This study was funded by the Stiftung Schleswig-Holsteinische Landschaft (Grant number: 2019/59). The sequencing costs were covered by the baseline funding from King Abdullah University of Science and Technology, Saudi Arabia, to Mark Tester.

## Supplementary Tables

**Supplementary Table 1.** Sampling stages and tissues from D-12082 and PI-614886 under two different photoperiod regimes. Samples were taken with two aims: RNA-sequencing (RNA-seq) and histological analysis. Plants were grown at 22°C under long-day conditions (LD, 16 h light) and short-day conditions (SD, 8 h light). Sequenced samples are shown in red.

**Supplementary Table 2.** Primer combinations used for RT-qPCR experiments.

**Supplementary Table 3.** Quality filtering and raw read statistics for the transcriptome libraries. Three biological replicates of shot apical meristem and leaf tissue from two accessions grown under two conditions were sequenced. Samples were collected weekly. Long-day conditions (LD): 16 h light, 22°C; short-day conditions (SD): 8 h light, 22°C. PE= paired-end.

**Supplementary Table 4.** The list of candidate genes putatively regulating flowering time in quinoa is mentioned in the text as having an associated function in quinoa and other species.

**Supplementary Table 5.** Gene Ontology (GO) annotation of differentially expressed genes (DEGs) between SD and LD in leaves of D-12082, which differentially respond to photoperiod to likely regulate flowering time (identifier A3 in Table 1 and Figure 1). The number of genes in each node of different GO hierarchy levels is shown for three GO categories, “Biological Process”, “Cellular Component” and “Molecular Function”.

**Supplementary Table 6.** KEGG classification of differentially expressed genes (DEGs) between SD and LD in leaves of D-12082, which differentially respond to photoperiod to likely regulate flowering time (identifier A3 in Table 1 and Figure 1).

**Supplementary Table 7.** Gene Ontology (GO) annotation of differentially expressed genes DEGs between PI-614886 and D-12082 at W2 and W3 under long-days (LD), which are putatively controlling flowering under LD (16 h light) (identifier D2 in Table 1 and Figure 1). The number of genes in each node of different GO hierarchy levels is shown for three GO categories, “Biological Process”, “Cellular Component” and “Molecular Function”.

**Supplementary Table 8.** KEGG classification of differentially expressed genes (DEGs) between PI-614886 and D-12082 at W2 and W3 under long-days (LD) in leaves, which are putatively controlling flowering under LD (16 h light) (identifier D2 in Table 1 and Figure 1).

**Supplementary Table 9.** Gene Ontology (GO) annotation of differentially expressed genes (DEGs) with similar spatial expression patterns in leaves between PI-614886 and D-12082, which likely regulate flowering time under short-days (8 h light) (identifier C3 in Table 1 and Figure 1). The number of genes in each node of different GO hierarchy levels is shown for three GO categories: “Biological Process”, “Cellular Component” and “Molecular Function”.

**Supplementary Table 10.** KEGG classification of differentially expressed genes (DEGs) with similar spatial expression patterns between PI-614886 and D-12082 under short-days (SD) in leaves, which likely regulate flowering time under SD (8 h light) (identifier C3 in Table 1 and Figure 1).

**Supplementary Table 11.** Gene Ontology (GO) annotation of differentially expressed genes (DEGs) in the shoot apical meristem (SAM) between PI-614886 and D-12082 at W2 and W3, which are likely involved in the regulation of the time to flower at the SAM (identifier E2 in Table 1 and Figure 1). The number of genes in each node of different GO hierarchy levels is shown for three GO categories: “Biological Process”, “Cellular Component” and “Molecular Function”.

**Supplementary Table 12.** KEGG classification of differentially expressed genes (DEGs) in the shoot apical meristem (SAM) between PI-614886 and D-12082 at W2 and W3, which are likely involved in the regulation of the time to flower at the SAM (identifier E2 in Table 1 and Figure 1).

**Supplementary Table 13.** Results of the blastx analysis for novel transcripts that are co-expressed with annotated DEGs that putatively regulate flowering in leaves and shoot apical meristem. The top 10% of novel transcripts with the highest degree and module membership (MM) > 0.7 from each module in the co-expression hierarchical clustering were used for blastx. Novel transcripts are those sequences assembled as transcripts by StringTie, which are not found as transcripts in the reference genome QQ74-V2.

**Supplementary Table 14.** Gene Ontology (GO) annotation of novel transcripts co-expressed with annotated DEGs that putatively regulate flowering in leaves and shoot apical meristem. The number of genes in each node of different GO hierarchy levels is shown for three GO categories: “Biological Process”, “Cellular Component” and “Molecular Function”. Novel transcripts are those sequences assembled as transcripts by StringTie, which are not found as transcripts in the reference genome QQ74-V2.

**Supplementary Table 15.** Novel transcripts in leaves and shoot apical meristem with sequence similarity to genes previously described in the literature as involved in flowering time or photoperiod regulation.

**Supplementary Table 16.** Candidate genes in common between this study and Maldonado-Taipe et al. (2022).

**Supplementary Table 17.** CONSTANS-like genes in the QQ74-V2 reference genome. Highlighted rows show the genes resulting as candidates after our transcriptome analysis.

## References

Adamczyk BJ, Lehti-Shiu MD, Fernandez DE. 2007. The MADS domain factors *AGL15* and *AGL18* act redundantly as repressors of the floral transition in Arabidopsis. The Plant Journal 50(6): 1007–1019.

Alandia G, Rodriguez J, Jacobsen S-E, Bazile D, Condori B. 2020. Global expansion of quinoa and challenges for the Andean region. Global Food Security 26: 100429.

Almeida-Silva F, Venancio TM. 2022. BioNERO: an all-in-one R/Bioconductor package for comprehensive and easy biological network reconstruction. Functional Integrative Genomics 22(1): 131–136.

Bäurle I, Dean C. 2006. The timing of developmental transitions in plants. Cell 125(4): 655–664.

Blázquez MA, Weigel D. 1999. Independent regulation of flowering by phytochrome B and gibberellins in Arabidopsis. Plant Physiology 120(4): 1025–1032.

Boyle EA, Li YI, Pritchard JK. 2017. An expanded view of complex traits: from polygenic to omnigenic. Cell 169(7): 1177–1186.

Buckler ES, Holland JB, Bradbury PJ, Acharya CB, Brown PJ, Browne C, Ersoz E, Flint-Garcia S, Garcia A, Glaubitz J. 2009. The genetic architecture of maize flowering time. Science 325(5941): 714–718.

Camejo D, Rodríguez P, Morales MA, Dell’Amico JM, Torrecillas A, Alarcón JJ. 2005. High temperature effects on photosynthetic activity of two tomato cultivars with different heat susceptibility. Journal of Plant Physiology 162(3): 281–289.

Cháb D, Kolář J, Olson MS, Štorchová H. 2008. Two *FLOWERING LOCUS T* (*FT)* homologs in *Chenopodium rubrum* differ in expression patterns. Planta 228(6): 929.

Chen L, Nan H, Kong L, Yue L, Yang H, Zhao Q, Fang C, Li H, Cheng Q, Lu S. 2020. Soybean *AP1* homologs control flowering time and plant height. Journal of Integrative Plant Biology 62(12): 1868–1879.

Chen Y, McCarthy D, Ritchie M, Robinson M, Smyth G, Hall E. 2020. edgeR: differential analysis of sequence read count data User’s Guide. Bioinformatics 26(1).

Cho LH, Yoon J, An G. 2017. The control of flowering time by environmental factors. The Plant Journal 90(4): 708–719.

Choi K, Kim J, Hwang H-J, Kim S, Park C, Kim SY, Lee I. 2011. The *FRIGIDA* complex activates transcription of *FLC*, a strong flowering repressor in Arabidopsis, by recruiting chromatin modification factors. The Plant Cell 23(1): 289–303.

Debernardi JM, Greenwood JR, Jean Finnegan E, Jernstedt J, Dubcovsky JJTPJ. 2020. APETALA 2-like genes AP2L2 and Q specify lemma identity and axillary floral meristem development in wheat. 101(1): 171–187.

Díaz-Valenzuela E, Hernández-Ríos D, Cibrián-Jaramillo A. 2023. The role of non-additive gene action on gene expression variation in plant domestication. EvoDevo 14(1): 1–14.

Drabešová J, Cháb D, Kolář J, Haškovcová K, Štorchová H. 2014. A dark–light transition triggers expression of the floral promoter *CrFTL1* and downregulates *CONSTANS*-like genes in a short-day plant *Chenopodium rubrum*. Journal of Experimental Botany 65(8): 2137–2146.

Dun EA, Ferguson BJ, Beveridge CA. 2006. Apical dominance and shoot branching. Divergent opinions or divergent mechanisms? Plant Physiology 142(3): 812–819.

Fuller HJ. 1949. Photoperiodic responses of *Chenopodium quinoa* Willd. and Amaranthus caudatus L. American Journal of Botany: 175–180.

Gaudinier A, Blackman BK. 2020. Evolutionary processes from the perspective of flowering time diversity. New Phytologist 225(5): 1883–1898.

Golicz AA, Steinfort U, Arya H, Singh MB, Bhalla PL. 2020. Analysis of the quinoa genome reveals conservation and divergence of the flowering pathways. Functional Integrative Genomics 20(2): 245–258.

Götz S, García-Gómez JM, Terol J, Williams TD, Nagaraj SH, Nueda MJ, Robles M, Talón M, Dopazo J, Conesa A. 2008. High-throughput functional annotation and data mining with the Blast2GO suite. Nucleic Acids Research 36(10): 3420–3435.

Granado-Rodríguez S, Aparicio N, Matías J, Pérez-Romero LF, Maestro I, Gracés I, Pedroche JJ, Haros CM, Fernandez-Garcia N, Del Hierro JN. 2021. Studying the impact of different field environmental conditions on seed quality of quinoa: The case of three different years changing seed nutritional traits in Southern Europe. Frontiers in plant science 12.

Gutierrez-Larruscain D, Krüger M, Abeyawardana OA, Belz C, Dobrev PI, Vaňková R, Eliášová K, Vondráková Z, Juříček M, Štorchová H. 2022. The transcriptomic (RNA-Sequencing) datasets collected in the course of floral induction in Chenopodium ficifolium 459. Data in Brief 43: 108333.

Higuchi Y, Hisamatsu T. 2015. *CsTFL1*, a constitutive local repressor of flowering, modulates floral initiation by antagonising florigen complex activity in chrysanthemum. Plant Science 237: 1–7.

Hu Y, Liang W, Yin C, Yang X, Ping B, Li A, Jia R, Chen M, Luo Z, Cai Q. 2015. Interactions of *OsMADS1* with floral homeotic genes in rice flower development. Molecular Plant 8(9): 1366–1384.

Iqbal S, Basra S, Saddiq MS, Yang A, Akhtar SS, Jacobsen S-E 2020. The extraordinary salt tolerance of quinoa. Emerging Research in Alternative Crops: Springer, 125–143.

Jarvis DE, Ho YS, Lightfoot DJ, Schmöckel SM, Li B, Borm TJ, Ohyanagi H, Mineta K, Michell CT, Saber N. 2017. The genome of *Chenopodium quinoa*. Nature 542(7641): 307.

Johanson U, West J, Lister C, Michaels S, Amasino R, Dean CJS. 2000. Molecular analysis of FRIGIDA, a major determinant of natural variation in Arabidopsis flowering time. 290(5490): 344–347.

Kanehisa M, Goto S. 2000. KEGG: kyoto encyclopedia of genes and genomes. Nucleic Acids Research 28(1): 27–30.

Kiani-Pouya A, Li L, Rasouli F, Zhang Z, Chen J, Yu M, Tahir A, Hedrich R, Shabala S, Zhang H. 2022. Transcriptome analyses of quinoa leaves revealed critical function of epidermal bladder cells in salt stress acclimation. Plant Stress 3: 100061.

Kim D, Langmead B, Salzberg SL. 2015. HISAT: a fast spliced aligner with low memory requirements. Nature Methods 12(4): 357–360.

Kim S-K, Yun C-H, Lee JH, Jang YH, Park H-Y, Kim J-K. 2008. *OsCO3*, a *CONSTANS-LIKE* gene, controls flowering by negatively regulating the expression of *FT-like* genes under SD conditions in rice. Planta 228(2): 355–365.

Komiya R, Ikegami A, Tamaki S, Yokoi S, Shimamoto K. 2008. Hd3a and RFT1 are essential for flowering in rice. Development 135(4).

Langfelder P, Horvath S. 2008. WGCNA: an R package for weighted correlation network analysis. BMC Bioinformatics 9(1): 1–13.

Langmead B, Salzberg S. 2012. Fast gapped-read alignment with Bowtie 2. Nature Methods 9(4): 357–359.

Lee J, Lee I. 2010. Regulation and function of *SOC1*, a flowering pathway integrator. Journal of Experimental Botany 61(9): 2247–2254.

Li B, Dewey C. 2011. RSEM: accurate transcript quantification from RNA-Seq data with or without a reference genome. BMC Bioinformatics 12: 1–16.

Liu L, Li C, Song S, Teo ZWN, Shen L, Wang Y, Jackson D, Yu H. 2018. *FTIP*-dependent *STM* trafficking regulates shoot meristem development in Arabidopsis. Cell Reports 23(6): 1879–1890.

Livak KJ, Schmittgen TD. 2001. Analysis of relative gene expression data using real-time quantitative PCR and the 2− ΔΔCT method. Methods 25(4): 402–408.

Lutz U, Pose D, Pfeifer M, Gundlach H, Hagmann J, Wang C, Weigel D, Mayer KF, Schmid M, Schwechheimer C. 2015. Modulation of ambient temperature-dependent flowering in Arabidopsis thaliana by natural variation of *FLOWERING LOCUS M*. PLoS Genetics 11(10): e1005588.

Maldonado-Taipe N, Barbier F, Schmid K, Jung C, Emrani N. 2022. High-Density Mapping of Quantitative Trait Loci Controlling Agronomically Important Traits in Quinoa (*Chenopodium quinoa* Willd.). Frontiers in plant science 13.

Maldonado-Taipe N, Patirange DS, Schmöckel SM, Jung C, Emrani N. 2021. Validation of suitable genes for normalization of diurnal gene expression studies in *Chenopodium quinoa*. PloS one 16(3): e0233821.

Más P, Kim W-Y, Somers DE, Kay SA. 2003. Targeted degradation of *TOC1* by *ZTL* modulates circadian function in *Arabidopsis thaliana*. Nature 426(6966): 567–570.

Mathieson I. 2021. The omnigenic model and polygenic prediction of complex traits. The American Journal of Human Genetics 108(9): 1558–1563.

Murai K, Miyamae M, Kato H, Takumi S, Ogihara Y. 2003. *WAP1*, a wheat *APETALA1* homolog, plays a central role in the phase transition from vegetative to reproductive growth. Plant Cell Physiology 44(12): 1255–1265.

Murphy KM, Matanguihan JB, Fuentes FF, Gómez-Pando LR, Jellen EN, Maughan PJ, Jarvis D. 2018. Quinoa breeding and genomics. Plant Breed. Rev. 42: 257–320.

Noh YS, Bizzell CM, Noh B, Schomburg FM, Amasino RM. 2004. *EARLY FLOWERING 5* acts as a floral repressor in Arabidopsis. The Plant Journal 38(4): 664–672.

Ogata T, Toyoshima M, Yamamizo-Oda C, Kobayashi Y, Fujii K, Tanaka K, Tanaka T, Mizukoshi H, Yasui Y, Nagatoshi Y. 2021. Virus-Mediated Transient Expression Techniques Enable Functional Genomics Studies and Modulations of Betalain Biosynthesis and Plant Height in Quinoa. Frontiers in plant science 12.

Patiranage DS, Asare E, Maldonado-Taipe N, Rey E, Emrani N, Tester M, Jung C. 2021. Haplotype variations of major flowering time genes in quinoa unveil their role in the adaptation to different environmental conditions. Plant, Cell & Environment.

Patiranage DSR, Rey E, Emrani N, Wellman G, Schmid K, Schmöckel SM, Tester M, Jung C. 2022. Genome-wide association study in quinoa reveals selection pattern typical for crops with a short breeding history. eLife 11: e66873.

Pertea G, Pertea M. 2020. GFF utilities: GffRead and GffCompare. Research 9: 304.

Pertea M, Kim D, Pertea GM, Leek JT, Salzberg SL. 2016. Transcript-level expression analysis of RNA-seq experiments with HISAT, StringTie and Ballgown. Nature Protocols 11(9): 1650–1667.

Pertea M, Pertea GM, Antonescu CM, Chang T-C, Mendell JT, Salzberg SL. 2015. StringTie enables improved reconstruction of a transcriptome from RNA-seq reads. Nature Biotechnology 33(3): 290–295.

Pin PA, Benlloch R, Bonnet D, Wremerth-Weich E, Kraft T, Gielen JJ, Nilsson O. 2010. An antagonistic pair of FT homologs mediates the control of flowering time in sugar beet. Science 330(6009): 1397–1400.

Putterill J, Varkonyi-Gasic E. 2016. FT and florigen long-distance flowering control in plants. Current Opinion in Plant Biology 33: 77–82.

Robinson MD, Oshlack A. 2010. A scaling normalization method for differential expression analysis of RNA-seq data. Genome Biology 11(3): 1–9.

Scortecci KC, Michaels SD, Amasino RM. 2001. Identification of a MADS-box gene, *FLOWERING LOCUS M*, that represses flowering. The Plant Journal 26(2): 229–236.

Shim Y, Lim C, Seong G, Choi Y, Kang K, Paek NC. 2022. The AP2/ERF transcription factor *LATE FLOWERING SEMI-DWARF* suppresses long-day-dependent repression of flowering. Plant, Cell & Environment 45(8): 2446–2459.

Song YH, Shim JS, Kinmonth-Schultz HA, Imaizumi T. 2015. Photoperiodic flowering: time measurement mechanisms in leaves. Annual Review of Plant Biology 66: 441–464.

Sosa-Zuniga V, Brito V, Fuentes F, Steinfort U. 2017. Phenological growth stages of quinoa (*Chenopodium quinoa*) based on the BBCH scale. Annals of Applied Biology 171(1): 117–124.

Sowiński P, Fronk J, Jończyk M, Grzybowski M, Kowalec P, Sobkowiak A. 2020. Maize response to low temperatures at the gene expression level: a critical survey of transcriptomic studies. Frontiers in plant science 11: 576941.

Stanschewski CS, Rey E, Fiene G, Craine EB, Wellman G, Melino VJ, SR Patiranage D, Johansen K, Schmöckel SM, Bertero D. 2021. Quinoa phenotyping methodologies: An international consensus. Plants 10(9): 1759.

Štorchová H, Drabešová J, Cháb D, Kolář J, Jellen EN. 2015. The introns in *FLOWERING LOCUS T-LIKE* (*FTL*) genes are useful markers for tracking paternity in tetraploid *Chenopodium quinoa* Willd. Genetic Resources and Crop Evolution 62: 913–925.

Suarez-Lopez P, Wheatley K, Robson F, Onouchi H, Valverde F, Coupland G. 2001. *CONSTANS* mediates between the circadian clock and the control of flowering in Arabidopsis. Nature 410(6832): 1116–1120.

Tautz D, Reeves G, Pallares LF. 2020. New experimental support for long standing concepts of polygenic genetics implies that the Mendelian genetic paradigm needs to be revised: The New (Old) Genetics, Version 1.0. NAL-live 2020(1): 1–15.

Tian H, Li Y, Wang C, Xu X, Zhang Y, Zeb Q, Zicola J, Fu Y, Turck F, Li L. 2021. Photoperiod-responsive changes in chromatin accessibility in phloem companion and epidermis cells of Arabidopsis leaves. The Plant Cell 33(3): 475–491.

Wang N, Qian W, Suppanz I, Wei L, Mao B, Long Y, Meng J, Müller AE, Jung C. 2011. Flowering time variation in oilseed rape (*Brassica napus* L.) is associated with allelic variation in the *FRIGIDA* homologue *BnaA. FRI. a*. Journal of Experimental Botany 62(15): 5641–5658.

Wang Q, Zuo Z, Wang X, Gu L, Yoshizumi T, Yang Z, Yang L, Liu Q, Liu W, Han Y-J. 2016. Photoactivation and inactivation of Arabidopsis cryptochrome 2. Science 354(6310): 343–347.

Wu M-F, Wagner D 2012. RNA in situ hybridization in Arabidopsis. RNA Abundance Analysis: Springer, 75–86.

Wu Q, Bai X, Luo Y, Li L, Nie M, Liu C, Ye X, Zou L, Xiang D. 2023. Identification of the global diurnal rhythmic transcripts, transcription factors and time-of-day specific cis elements in *Chenopodium quinoa*. BMC Plant Biology 23(1): 1–19.

Wu Q, Luo Y, Wu X, Bai X, Ye X, Liu C, Wan Y, Xiang D, Li Q, Zou L. 2021. Identification of the specific long-noncoding RNAs involved in night-break mediated flowering retardation in *Chenopodium quinoa*. BMC genomics 22(1): 1–18.

Yan L, Loukoianov A, Tranquilli G, Helguera M, Fahima T, Dubcovsky J. 2003. Positional cloning of the wheat vernalization gene *VRN1*. Proceedings of the National Academy of Sciences 100(10): 6263–6268.

Zheng L, Zhao Y, Gan Y, Li H, Luo S, Liu X, Li Y, Shao Q, Zhang H, Zhao Y. 2022. Full-Length Transcriptome Sequencing Reveals the Impact of Cold Stress on Alternative Splicing in Quinoa. International journal of molecular sciences 23(10): 5724.

