## Supplementary Figures for "Leaf and shoot apical meristem transcriptomes of quinoa (*Chenopodium quinoa* Willd.) in response to photoperiod and plant development"

Supplementary Figure 1

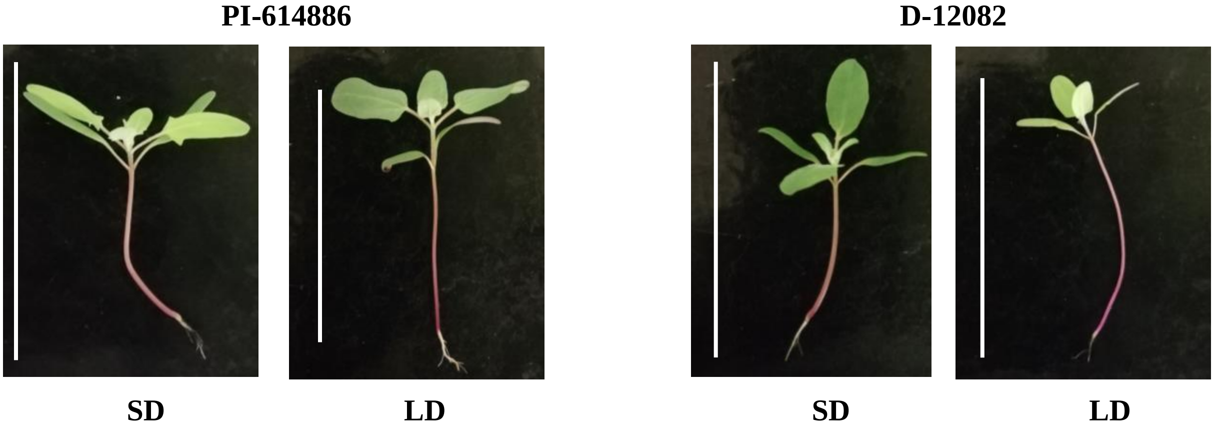

Supplementary Figure 2

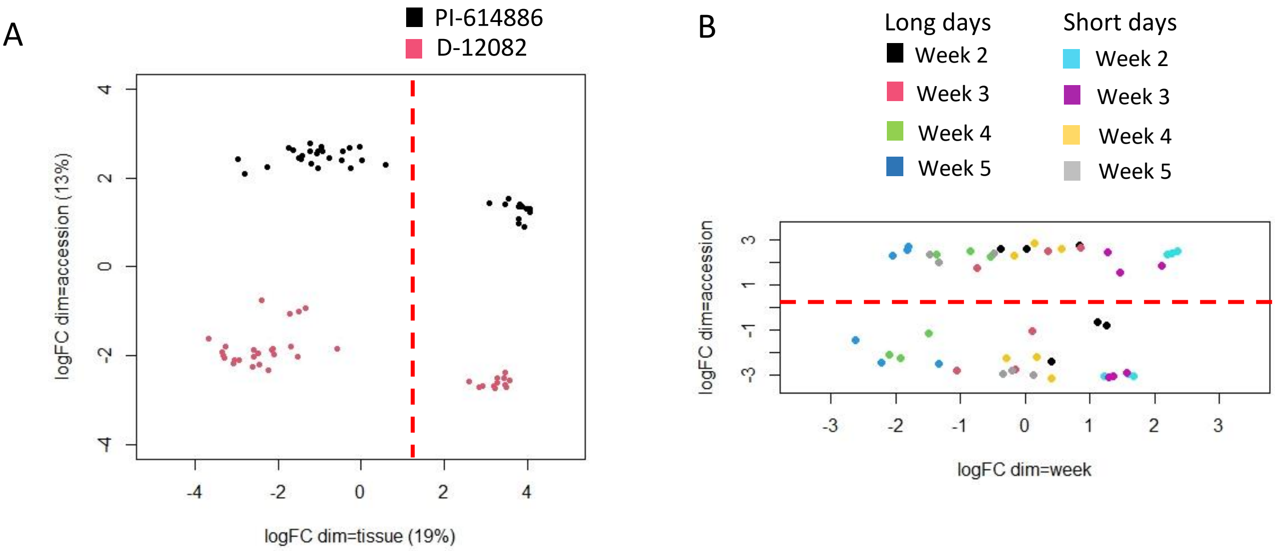

Supplementary Figure 3

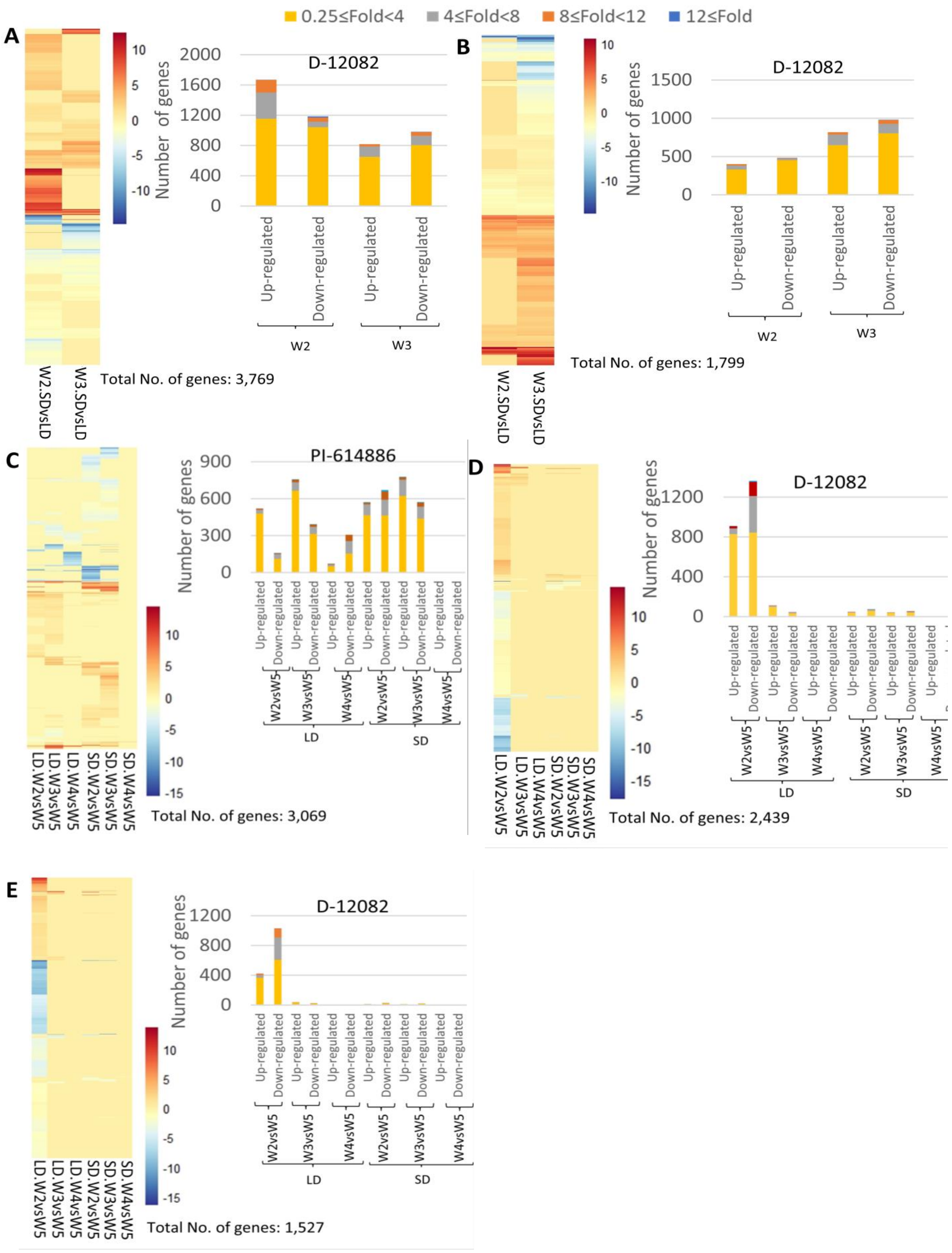

Supplementary Figure 4

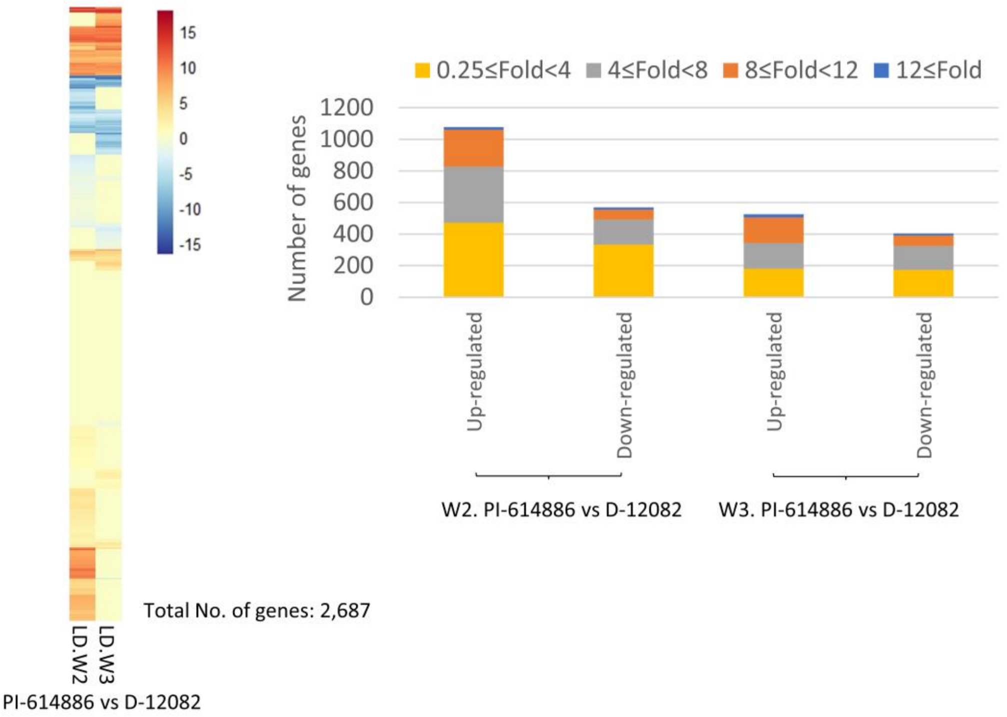

Supplementary Figure 5

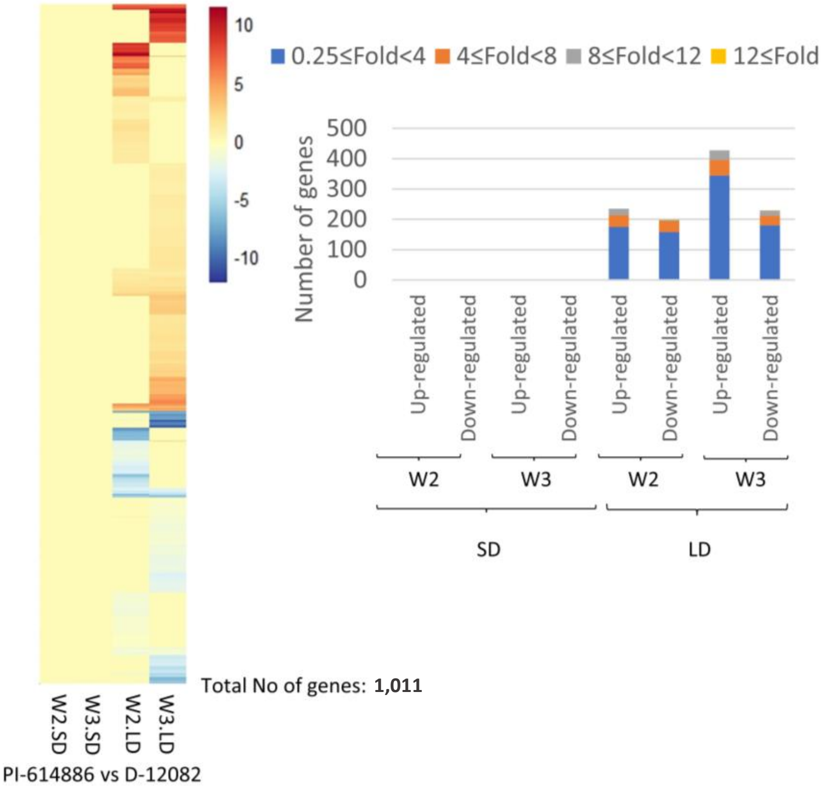

Supplementary Figure 6

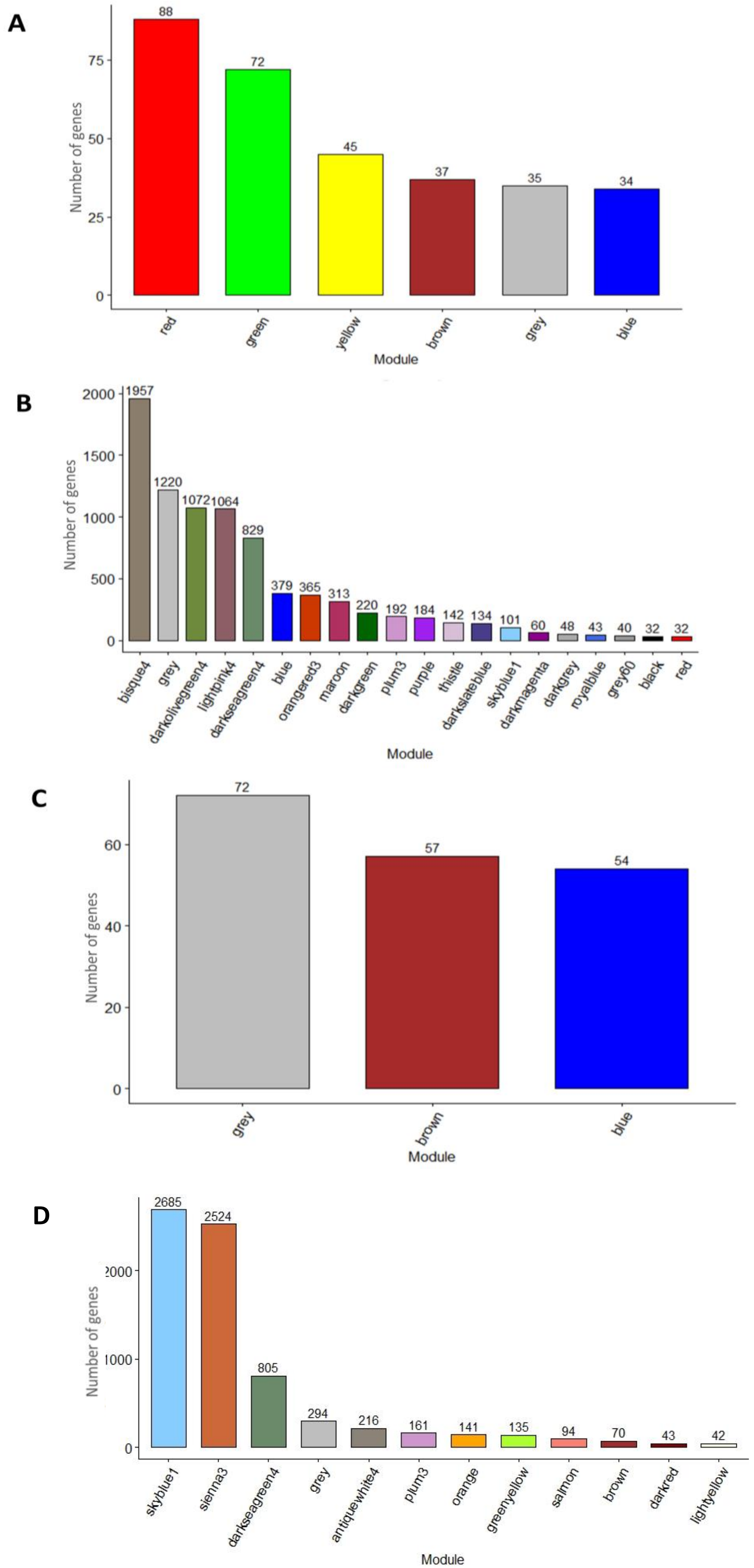

Supplementary Figure 7

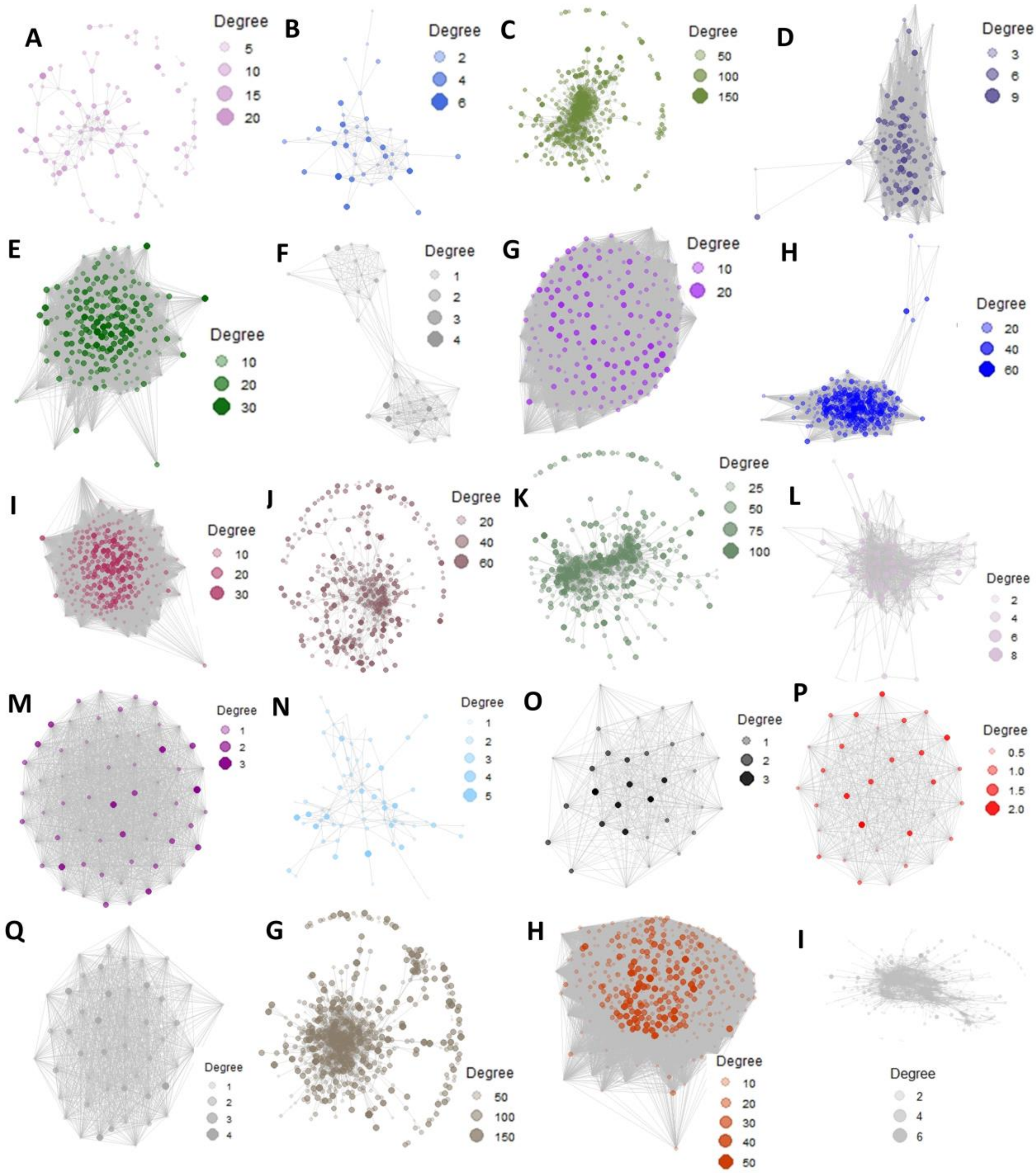

### Supplementary Figure 8

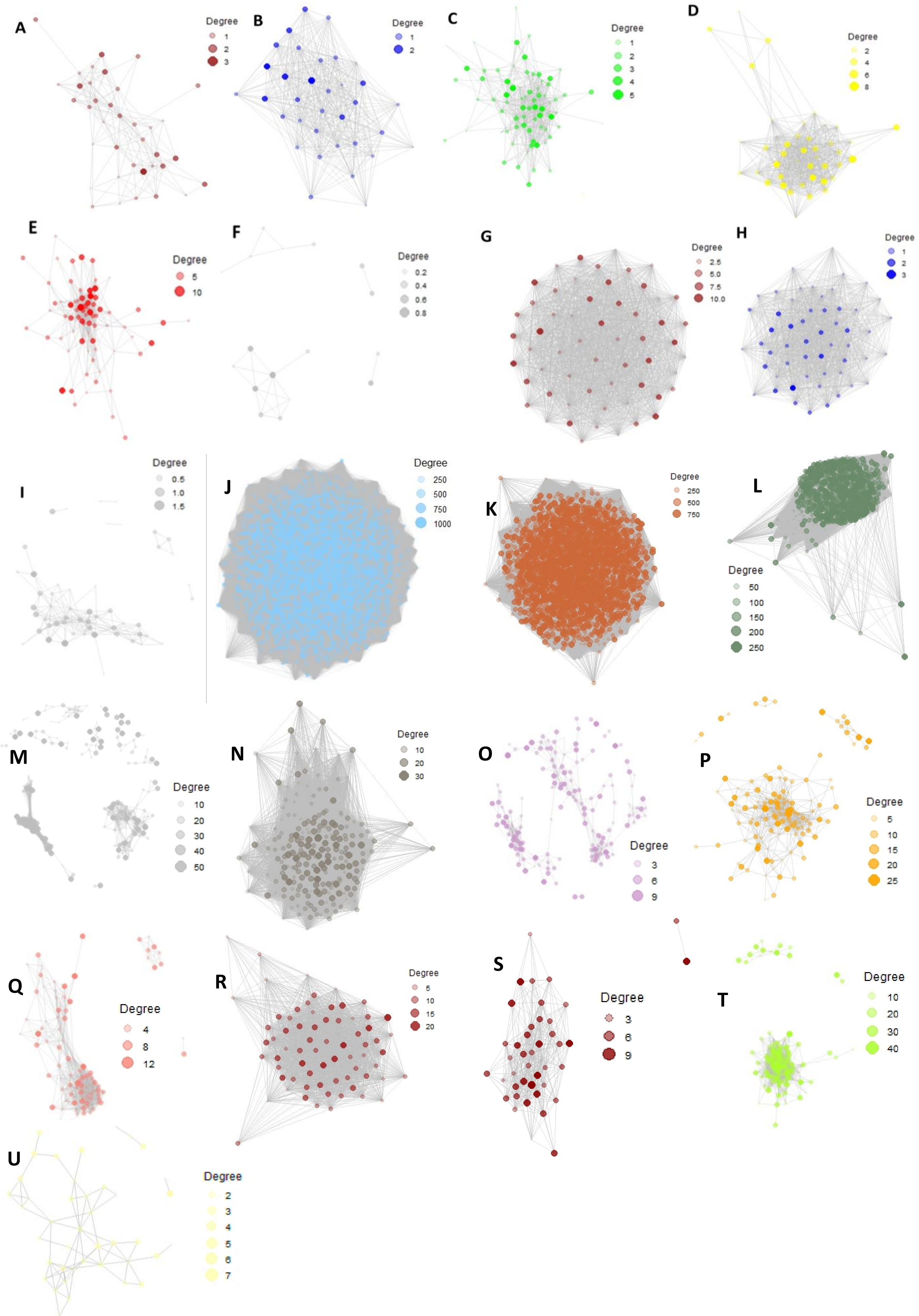

Supplementary Figure 9

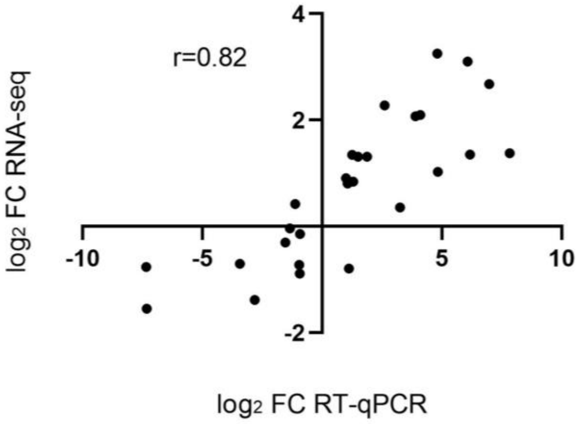

Supplementary Figure 10

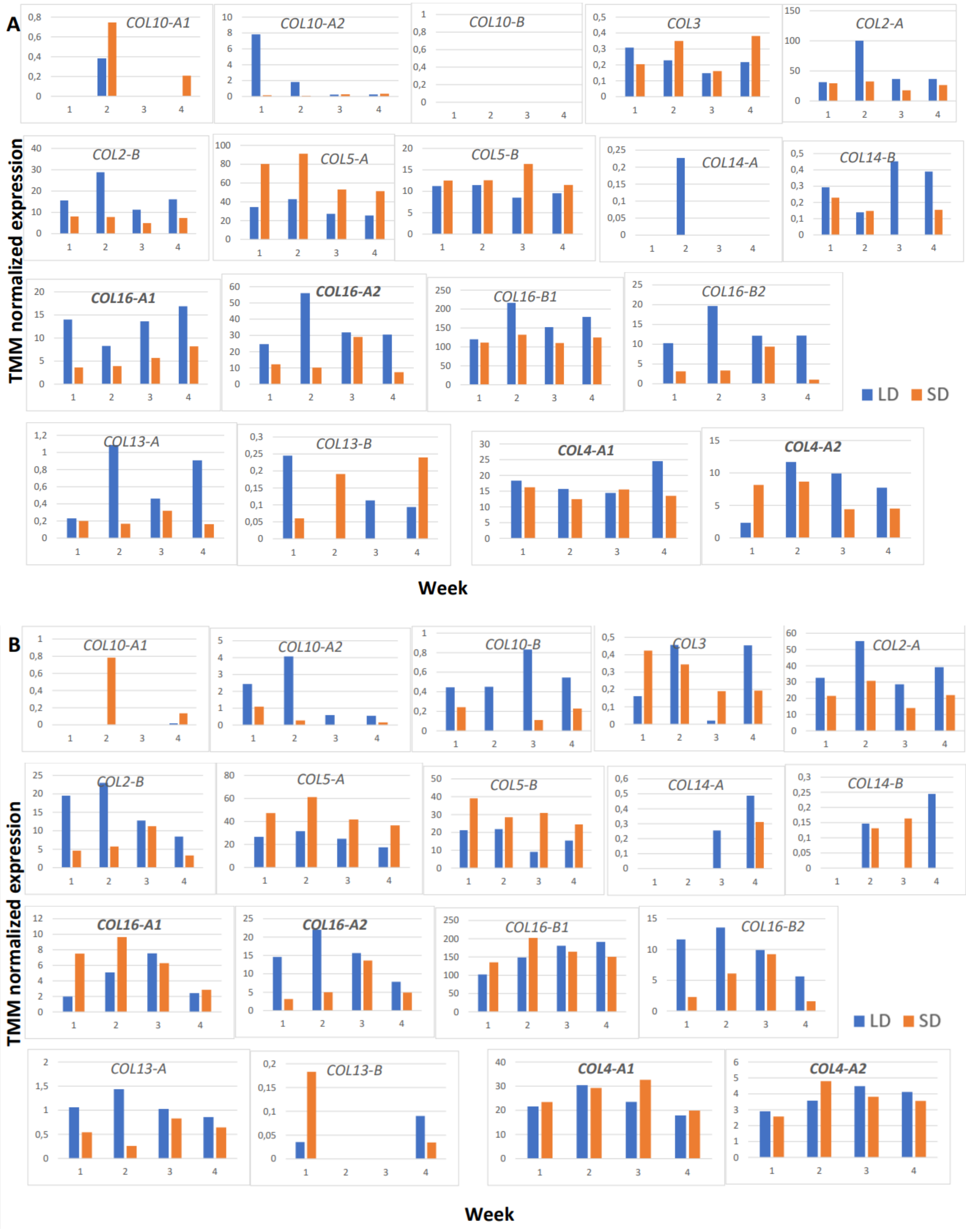
